# Omicron-specific mRNA vaccination alone and as a heterologous booster against SARS-CoV-2

**DOI:** 10.1101/2022.02.14.480449

**Authors:** Zhenhao Fang, Lei Peng, Renata Filler, Kazushi Suzuki, Andrew McNamara, Qianqian Lin, Paul A. Renauer, Luojia Yang, Bridget Menasche, Angie Sanchez, Ping Ren, Qiancheng Xiong, Madison Strine, Paul Clark, Chenxiang Lin, Albert I. Ko, Nathan D. Grubaugh, Craig B. Wilen, Sidi Chen

## Abstract

The Omicron variant of severe acute respiratory syndrome coronavirus 2 (SARS-CoV-2) has high transmissibility and recently swept the globe. Due to the extensive number of mutations, this variant has high level of immune evasion, which drastically reduced the efficacy of existing antibodies and vaccines. Thus, it is important to test an Omicron-specific vaccine, evaluate its immune response against Omicron and other variants, and compare its immunogenicity as boosters with existing vaccine designed against the reference wildtype virus (WT). Here, we generated an Omicron-specific lipid nanoparticle (LNP) mRNA vaccine candidate, and tested its activity in animals, both alone and as a heterologous booster to existing WT mRNA vaccine. Our Omicron-specific LNP-mRNA vaccine elicited strong and specific antibody response in vaccination-naïve mice. Mice that received two-dose WT LNP-mRNA, the one mimicking the commonly used Pfizer/Moderna mRNA vaccine, showed a >40-fold reduction in neutralization potency against Omicron variant than that against WT two weeks post second dose, which further reduced to background level >3 months post second dose. As a booster shot for two-dose WT mRNA vaccinated mice, a single dose of either a homologous booster with WT LNP-mRNA or a heterologous booster with Omicron LNP-mRNA restored the waning antibody response against Omicron, with over 40-fold increase at two weeks post injection as compared to right before booster. Interestingly, the heterologous Omicron LNP-mRNA booster elicited neutralizing titers 10-20 fold higher than the homologous WT booster against the Omicron variant, with comparable titers against the Delta variant. All three types of vaccination, including Omicron mRNA alone, WT mRNA homologous booster, and Omicron heterologous booster, elicited broad binding antibody responses against SARS-CoV-2 WA-1, Beta, and Delta variants, as well as other *Betacoronavirus* species such as SARS-CoV, but not Middle East respiratory syndrome coronavirus (MERS-CoV). These data provided direct proof-of-concept assessments of an Omicron-specific mRNA vaccination *in vivo*, both alone and as a heterologous booster to the existing widely-used WT mRNA vaccine form.

## Introduction

Since its first identification in specimen collected in November 2021^1^, the Omicron variant (lineage B.1.1.529) of SARS-CoV-2 has rapidly spread across the globe^2–5^. The Omicron variant was associated with increased risk of reinfection according to a population-level evidence in South Africa^6^. Two days after its initial report to World Health Organization (WHO), Omicron was designated as a variant of concern (VoC) by WHO on November 26^7^. Population-level data indicated that Omicron has become the dominant variant in South Africa in mid-November^8^, only one week after the first traceable case. Similarly in January 2022, Omicron variant has dominated newly diagnosed cases in many states of the US^9^, Canada^10^ and UK^4^. Omicron variant drove the fourth “wave” of Coronavirus Disease 2019 (COVID-19) in South Africa^11^ and around the world^12^. Its case doubling time, every 3-4 days, is faster than previous waves^12^, hinting its increased intrinsic transmissibility and/or immune evasion. A number of urgent questions on Omicron have quickly become central concerns, which include whether and when Omicron-specific vaccines or therapeutic antibodies will be effective against Omicron variant.

Mounting clinical and laboratory evidence have shown that most therapeutic and natural antibodies for COVID-19 failed to retain potency against Omicron^13–18^. The Omicron variant has 60 mutations compared to the ancestral variant’s reference sequence (also referred to as prototypic virus / variant, reference, wildtype (WT), Wuhan-Hu-1, or Wuhan-1, in lineage A)^5^. There are 50 nonsynonymous, 8 synonymous, and 2 non-coding mutations in Omicron, of which many are not observed in any other variants. There are a total of 32 mutations in the spike gene, which encodes the main antigen target of therapeutic antibodies and of many widely administered vaccines. This results in 30 amino acid changes, three small deletions, and one small insertion, of which 15 are within the receptor-binding domain (RBD)^19^. Due to its extensive number of mutations, this variant has high level of immune evasion, which drastically reduced the efficacy of existing antibodies and vaccines. Omicron spike mutations are concerning as they cluster on known neutralizing antibody epitopes^15^ and some of them have well-characterized consequences such as immune evasion and higher infectivity. In fact, recent reports showed that the majority of the existing monoclonal antibodies developed against SARS-CoV-2 have dramatic reduction in effectiveness against the Omicron variant^13–18^, leading to recall or exclusion of recommended use of certain therapeutic antibodies under emergency authorization^20^.

The mRNA vaccines have achieved immense success in curbing the viral spread and reducing the risk of hospitalization and death of COVID-19^21,22^. However, a significant drop of mRNA vaccine’s effectiveness against Omicron has been reported from clinical^8^ and laboratory studies of samples of vaccinated individuals^23,24^. In light of the heavily altered antigen landscape of Omicron spike, assessing the efficacy of Omicron-specific mRNA vaccine is urgently needed. A number of critical questions regarding Omicron-specific mRNA vaccine need to be addressed. For examples: What is the immunogenicity of Omicron spike used in a vaccine form? Whether potent antibody immunity can be induced by an Omicron-specific mRNA vaccine, and how well does that neutralize the Omicron variant? If, and how, does the immune response induced by the Omicron-specific react to other variants, such as WA-1 (lineage A with a spike gene identical to WT or Wuhan-1) or Delta (lineage B.1.617.2)? Because a large share of world population received authorized Pfizer/BioNTech or Moderna mRNA vaccines that encode the reference (WT) spike antigen, it is important to know whether an Omicron mRNA vaccine can boost the waning immunity of existing vaccinated population. Clinical data showed that heterologous boosting with different types of COVID-19 vaccines elicited neutralizing titers similar to or greater than homologous boosting^25,26^. It is critical to compare the immunogenicity and efficacy of a heterologous Omicron booster with a homologous WT booster. Last but not least, as the antibody epitopes are closely related to their cross reactivity and susceptibility to variant mutations, it is crucial to know if and to what extent WT or Omicron mRNA vaccine can elicit plasma antibodies possessing broad antibody responses to SARS-CoV-2 variants and *Betacoronavirus* species.

To answer some of these questions, we directly generated an Omicron-specific lipid nanoparticle (LNP) mRNA vaccine candidate that encodes an engineered full-length Omicron spike with HexaPro mutations, and evaluated its effect alone, and compared its immunogenicity with WT LNP-mRNA as booster shots after SARS-CoV-2 WT mRNA vaccination in animal models.

## Results

### Design, generation and physical characterization of an Omicron-specific LNP-mRNA vaccine candidate

We designed an Omicron-specific LNP-mRNA vaccine candidate based on the full-length spike sequence of the Omicron variant (lineage B.1.1.529/BA.1) from two North America patients identified on Nov23^rd^, 2021 (GISAID EpiCoV: EPI_ISL_6826713 and EPI_ISL_6826714). The spike coding sequence of Wuhan-Hu-1 (WT) and Omicron variant were flanked by 5’ UTR, 3’ UTR and 3’ PolyA tail (**Figure 1A**). We introduced six Proline mutations (HexaPro) to the spike gene sequence, as they were reported to improve spike protein stability and prefusion state ^27^. The furin cleave site (RRAR) in spike was replaced with GSAS stretch to keep integrity of S1 and S2 units. We then encapsulated the transcribed spike mRNA into lipid nanoparticles to produce WT and Omicron LNP-mRNAs, and characterized the quality and biophysical properties by downstream assays including dynamic light scattering, transmission electron microscope and receptor binding assay.

**Figure 1.**
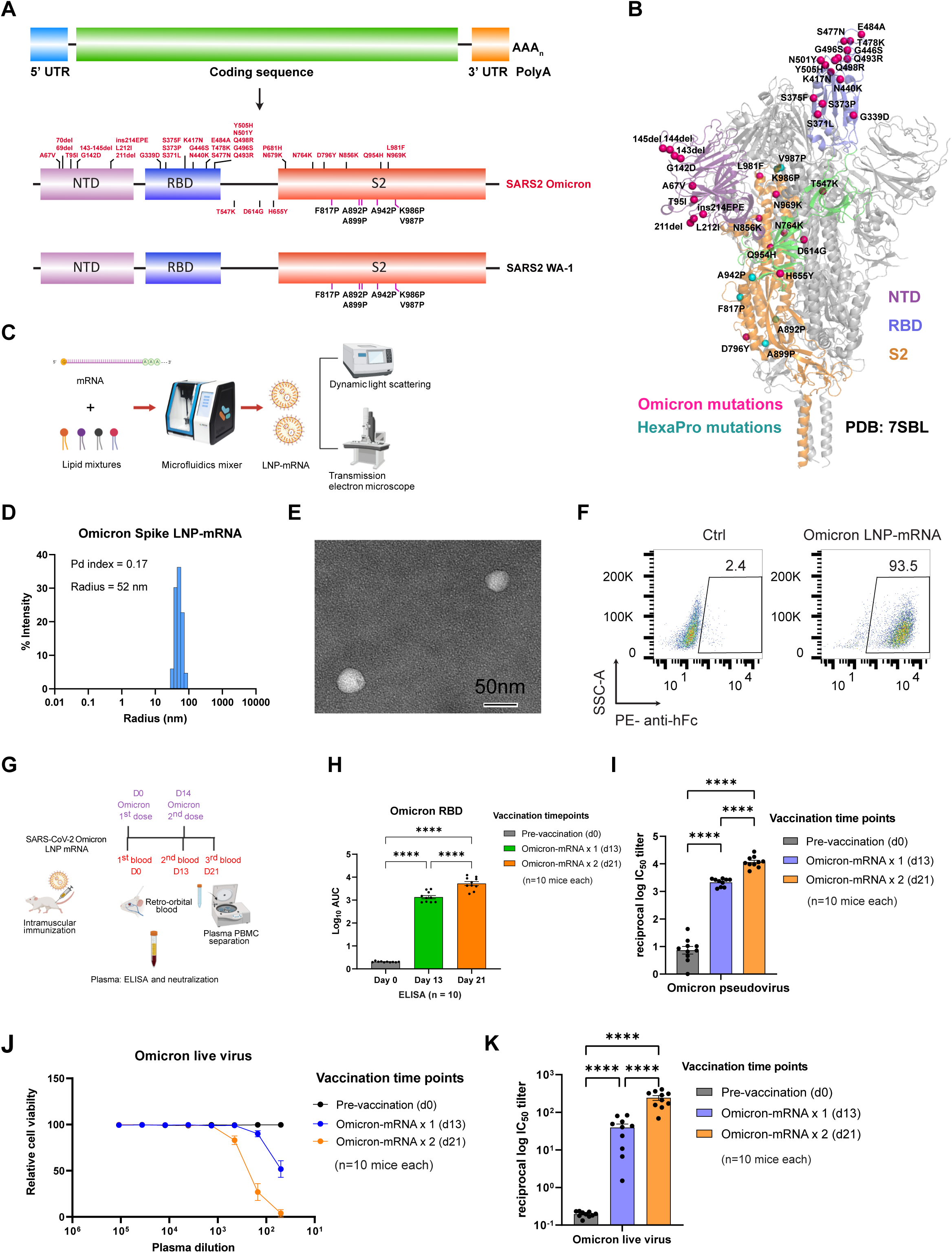
Omicron-specific LNP-mRNA vaccine elicited neutralizing antibodies against SARS-CoV-2 Omicron variant. **A**, Illustration of mRNA vaccine construct expressing SARS-CoV-2 WT and Omicron spike genes. The spike open reading frame were flanked by 5’ untranslated region (UTR), 3’ UTR and polyA tail. The Omicron mutations (red) and HexaPro mutations (black) were numbered based on WA-1 spike residue number. **B**, Distribution of Omicron spike mutations (magenta) were displayed in one protomer of spike trimer of which NTD, RBD, hinge region and S2 were colored in purple, blue, green and orange respectively (PDB: 7SBL). The HexaPro mutations in S2 were colored in cyan. **C**, Schematics illustrating the formulation and biophysical characterization of LNP-mRNA. **D**, Dynamic light scattering derived histogram depicting the particle radius distribution of Omicron spike LNP-mRNA. **E**, Omicron LNP-mRNA image collected on transmission electron microscope. **F**, human ACE2 receptor binding of LNP-mRNA encoding Omicron spike expressed in 293T cells as detected by human ACE2-Fc fusion protein and PE-anti-human Fc antibody on Flow cytometry. **G**, Immunization and sample collection schedule. Retro-orbital blood were collected prior Omicron LNP-mRNA vaccination on day 0, day 13 and day 21. Ten mice (n=10) were intramuscularly injected with 10 µg Omicron LNP-mRNA on day 0 (prime, Omicron x 1) and day 14 (boost, Omicron x 2). The plasma and peripheral blood mononuclear cells (PBMCs) were separated from blood for downstream assays. The slight offset of the labels reflects the fact that each of the blood collections were perform prior to the vaccination injections. Data were collected from two independent experiments and each experiment has five mice. **H**, Binding antibody titers of plasma from mice vaccinated with Omicron LNP-mRNA against Omicron spike RBD as quantified by area under curve of log_10_-transformed titration curve (Log_10_ AUC) in Figure S1. Each dot in bar graphs represents value from one mouse (n = 10 mice). **I**, Neutralization of Omicron pseudovirus by plasma from Omicron LNP-mRNA vaccinated mice. **J**, Omicron live virus titration curves over serial dilution points of plasma from mice before and after immunization with Omicron LNP-mRNA at defined time points. Data of each sample were collected from three replicates (n = 10 mice). **K**, Neutralization of Omicron infectious virus by plasma from Omicron LNP-mRNA vaccinated mice (n = 10 mice). Data on dot-bar plots are shown as mean ± s.e.m. with individual data points in plots. One-way ANOVA with Tukey’s multiple comparisons test was used to assess statistical significance. Statistical significance labels: * p < 0.05; ** p < 0.01; *** p < 0.001; **** p < 0.0001. Non-significant comparisons are not shown, unless otherwise noted as n.s., not significant.

The dynamic light scattering and transmission electron microscope were applied to evaluate the size distribution and shape of Omicron LNP-mRNA, which showed a monodispersed sphere shape with an average radius of 52 nm and polydispersity index of 0.17 (**Figure 1C-1E**). To evaluate the effectiveness of LNP-mRNA mediated Omicron spike expression in cells as well as the receptor binding ability of the designed Omicron HexaPro spike, Omicron LNP-mRNA was directly added to HEK293T cells 16 hours before subjecting cells to flow cytometry. Evident surface expression of functional Omicron HexaPro spike capable of binding to human angiotensin-converting enzyme-2 (hACE2) was observed by staining cells with hACE2-Fc fusion protein and PE anti-Fc secondary antibody (**Figure 1F**). These data showed that the Omicron spike sequence was successfully encoded into an mRNA, encapsulated into the LNP, can be introduced into mammalian cells efficiently without additional manipulation, and express functional spike protein that binds to hACE2.

### Specific binding and neutralizing antibody response elicited by Omicron LNP-mRNA against the Omicron variant

After ensuring functional spike expression mediated by Omicron LNP-mRNA, we proceeded to characterizing the immunogenicity of Omicron LNP-mRNA *in vivo*. In order to test rapid immune elicitation against Omicron variant, we performed the following vaccination and testing schedule. Two doses of 10 µg Omicron LNP-mRNA, as prime and boost two weeks apart were intramuscularly injected into ten C57BL/6Ncr (B6) mice (**Figure 1G; Figure S1A**). Retro-orbital blood was collected prior to immunization on day 0, 13 and 21, i.e. two weeks post prime (one day before boost), and one week post boost. We then isolated plasma from blood, which was used in enzyme-linked immunosorbent assay (ELISA) and neutralization assay to quantify binding and neutralizing antibody titers. A significant increase in antibody titers against Omicron spike RBD was observed in ELISA and neutralization assays from plasma samples post prime and boost (**Figure 1H-1I; Figure S1A-B**). We performed neutralization with infectious virus (also commonly referred to as authentic virus or live virus) using a local SARS-CoV-2 Omicron isolate in a biosafety level 3 (BSL3) setting (Methods), and validated that the plasma samples from mice vaccinated with Omicron-specific LNP-mRNA showed potent neutralization activity against infectious Omicron virus, with significant prime / boost effect (**Figure 1J-K**). These data showed that the Omicron LNP-mRNA induced strong and specific antibody responses in vaccinated mice.

### Waning immunity of WT LNP-mRNA immunized mice

In light of the wide coverage of the ancestral WT-based LNP-mRNA vaccine (to model those widely administered in the current general population), we sought to test: (i) the effect of WT LNP-mRNA vaccination against Omicron variant, (ii) the decay of immunity induced by WT LNP-mRNA over time, and (iii) whether a homologous WT LNP-mRNA booster or a heterologous Omicron LNP-mRNA booster could enhance the waning immunity against Omicron variant, WA-1 and/or Delta variant, and if there is a difference between homologous and heterologous boost. To gain initial answers to these questions in animal models, we sequentially vaccinated two cohorts of B6 mice with two doses of WT and one dose of WT or Omicron LNP-mRNA booster in two independent experiments (Batch 1 in **Figure S2** and batch 2 in **Figure S3**). Over 100-day interval between 2^nd^ dose of WT and WT/Omicron booster was ensured in order to observe the waning immunity in WT-vaccinated mice (the combined and individual datasets from the two independent experiments were presented in **Figures 2** and **S2-S3** respectively). We collected blood samples of these animals in a rational time series, including day 35 (2 weeks post 2^nd^ dose of WT LNP-mRNA), >3.5 months post 2^nd^ doses of WT LNP-mRNA (day 127 in batch 1 or day 166 in batch 2, immediately before WT/Omicron booster), ∼2 weeks post WT/Omicron LNP-mRNA booster (day 140, one day before the second Omicron booster in batch 1 or day 180 in batch 2), and day 148 (1 week post two doses of Omicron LNP-mRNA vaccination in batch 1).

**Figure 2.**
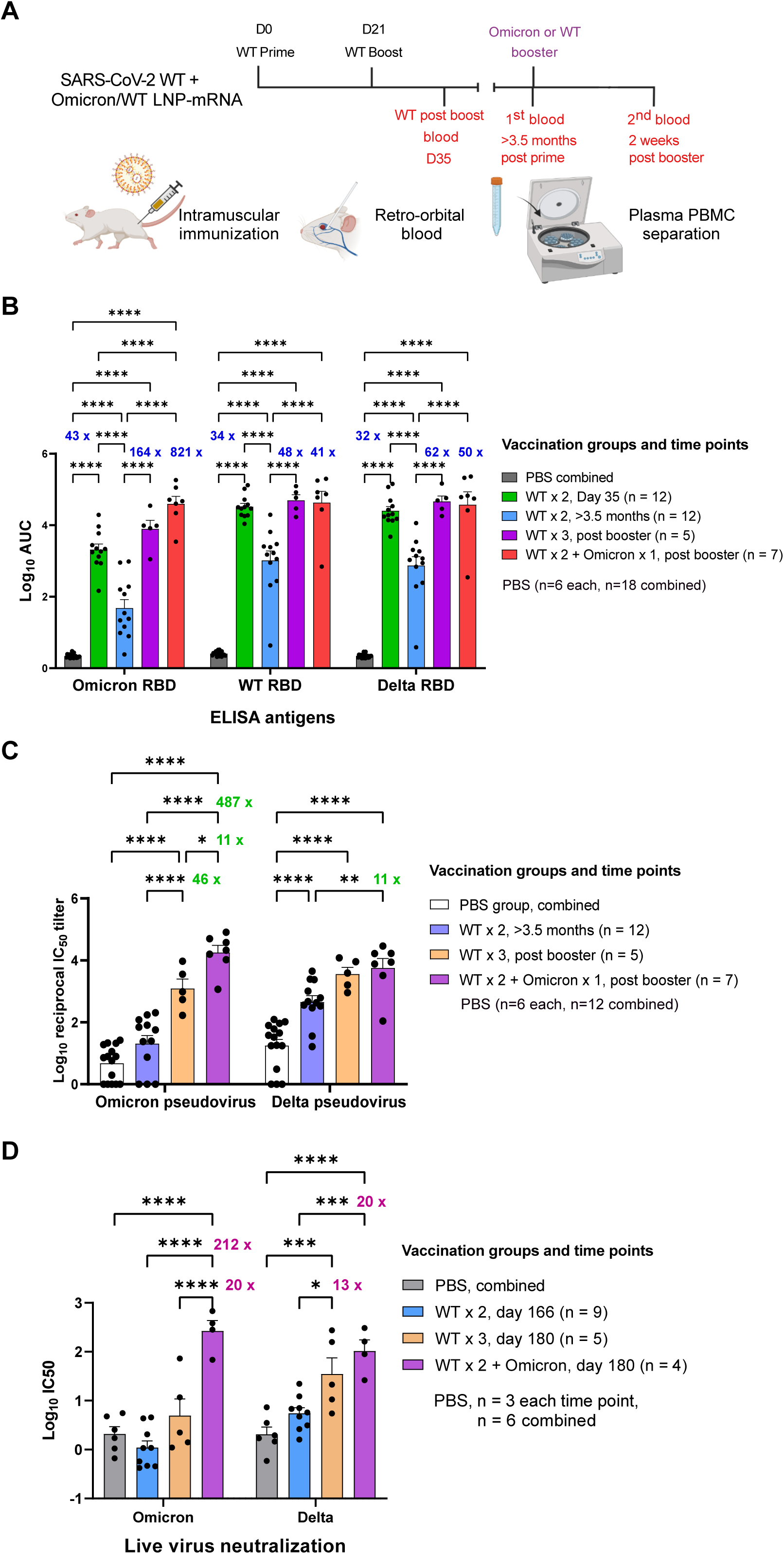
Heterologous booster with Omicron LNP-mRNA as compared to homologous booster with WT LNP-mRNA in mice that previously received a two-dose WT LNP-mRNA vaccination. **A**, Schematics showing the immunization and blood sampling schedule of mice administered with 1 µg WT LNP-mRNA prime (WT x 1) and boost (WT x 2) as well as 10 µg WT or Omicron-specific LNP-mRNA booster shots. The data was collected and combined from two independent experiments shown in Extended Data Figures S2 and S3. **B**, Bar graph comparing binding antibody titers of mice administered with PBS or WT and Omicron LNP-mRNA against Omicron, Delta and WA-1 RBD (ELISA antigens). The antibody titers were quantified as Log_10_ AUC based on titration curves in Extended Data Figure 1A. PBS sub-groups (n=6 each) collected from different matched time points showed no statistical differences between each other, and were combined as one group (n=18). **C**, Pseudovirus neutralizing antibody titers in the form of log_10_-transformed reciprocal IC50 calculated from fitting the titration curve with a logistic regression model (n = 12 before booster, n=5 in WT x 3, n = 7 in WT x 2 + Omicron). **D,** Infectious virus neutralization titer comparisons between mice before and after vaccination with WT or Omicron boosters (n = 9 before booster, n=5 in WT x 3, n = 4 in WT x 2 + Omicron). Titer ratios were indicated in each graph and fold change described in manuscript is calculated from (ratio - 1). Data on dot-bar plots are shown as mean ± s.e.m. with individual data points in plots. Two-way ANOVA with Tukey’s multiple comparisons test was used to assess statistical significance. Statistical significance labels: * p < 0.05; ** p < 0.01; *** p < 0.001; **** p < 0.0001. Non-significant comparisons are not shown, unless otherwise noted as n.s., not significant.

Plasma samples were isolated from blood samples and analyzed in ELISA and neutralization assays against SARS-CoV-2 Omicron, Delta or WA-1. Comparing to the titers against WA-1 and Delta RBD, the binding antibody titers against Omicron RBD elicited by WT mRNA-LNP were significantly weaker in samples from both day 35 and >3.5 months (**Figures 2B, S4 and S5**). The group average Omicron reactivity is 15-fold (day 35) and 21-fold (>3.5 months) lower than that of WT RBD (fold change = ratio -1), and 11-fold (day 35) and 14-fold (>3.5 months) lower than Delta (**Figure S5**). A steep (orders of magnitude) drop of antibody titers from mice immunized with WT LNP-mRNA was observed after three months (day 35 vs. >3.5 months) from all three RBD datasets. It is worth noting that the antibody titers >3.5 months post WT boost decreased to a level that is near-baseline (Phosphate buffered saline, PBS controls, **Figure 2**), particularly for titers against Omicron RBD.

### Heterologous booster with Omicron LNP-mRNA as compared to homologous booster with WT LNP-mRNA in mice that previously received a two-dose WT LNP-mRNA vaccination

A single dose booster shot, either a homologous booster with WT LNP-mRNA, or a heterologous booster with Omicron LNP-mRNA, drastically increased the antibody titers against Omicron RBD, by over 100-fold as compared to the sample right before booster shot (**Figure 2B**), reaching a level comparable to the post-boost titer by Omicron LNP-mRNA alone (**Figure 1H**). The mice that received the Omicron LNP-mRNA booster showed a trend of higher binding antibody titer against Omicron RBD than those administered with WT booster. Interestingly, the Omicron LNP-mRNA shot boosted not only titers against Omicron RBD, but also titers against Delta and WA-1 RBD, of which levels were comparable with those elicited by WT LNP-mRNA booster (**Figure 2B**). For both WT and Omicron boosters, the extent of titer increase was more drastic in the Omicron RBD dataset than other RBD datasets, signifying the extra benefit of booster shots against Omicron variant (**Figure 2B**). The antibody titers did not increase one week after a second booster of Omicron LNP-mRNA (**Figure S2B**).

Because pseudovirus neutralization is a relatively safer and widely-used assay that strongly correlates with infectious virus results and has been regarded as a standard proxy by the field ^24,28–30^, we set out to first use pseudovirus neutralization assay to measure the neutralizing antibody responses induced by Omicron LNP-mRNA booster in these animals. We first generated human immunodeficiency virus-1 (HIV-1) based Omicron pseudovirus system, which contains identical Omicron mutations in vaccine antigen, but lacks the HexaPro or furin site modifications. Interestingly, we found that under exactly the same virus production and assay conditions, the Omicron pseudovirus has higher infectivity than both WA-1 (8x increase) and Delta (4x) pseudoviruses (**Figure S6A-C**), which was also observed by another group^24^, in concordance with the Omicron - hACE2 interactions from biophysical and structural studies ^31,32^, and correlated with higher transmissibility reported previously^24,33,34^.

We then normalized the pseudoviruses by functional titers (number of infected cells / volume), and used this system to perform pseudovirus neutralization assays on all of plasma samples collected (**Figure S6D-E**). The neutralization results showed a consistent overall pattern as ELISA results, with a stronger contrast among titers against Omicron pseudovirus (**Figure 2C**). On day 35 and > 3.5 months post WT boost, the mice showed significantly lower neutralizing antibody titers against Omicron variant than titers against Delta variant or WA-1 (**Figure S7A-B**). For the samples two weeks post boost (day 35), the group average Omicron neutralization reactivity is 40-fold lower than that of WA-1 RBD, and 10-fold lower than Delta (**Figure S7B**). When comparing samples collected on day 35 and >3.5 months post WT boost, around two orders of magnitude (10s∼100s of fold change) time-dependent titer reduction was unequivocally observed in all three pseudovirus neutralization data (**Figure S2C**). The Omicron-neutralization activity of WT vaccinated mice >3.5 months post boost was as low as PBS background (**Figure 2C**). These data suggested that there was waning antibody immunity in the standard two-dose WT vaccinated animals, which lost neutralization ability against the Omicron variant pseudovirus.

A single booster shot of WT or Omicron LNP-mRNA vaccine enhanced the antibody titers against Omicron variant two weeks after the injection by >40-fold (**Figure 2C**). The heterologous Omicron LNP-mRNA booster induced significantly higher neutralizing titer against Omicron pseudovirus than the homologous WT LNP-mRNA booster (**Figure 2C**). The neutralizing titer after this surge by Omicron vaccine numerically surpassed the titer two weeks post WT vaccine boost (day 35, **Figure S2C**). Interestingly, the Omicron mRNA vaccine also rescued the antibody titers against Delta and WA-1 pseudoviruses, with two orders of magnitude increase in both ELISA titers and neutralization activity (**Figure 2B-2C**). The neutralization titers of Delta pseudovirus were found similar between WT and Omicron booster groups (**Figure 2C**). A second booster shot two weeks after the first of Omicron mRNA vaccine yielded little increase in neutralization activity against Omicron, WA-1 or Delta variants at the time measured (day 148, 1 week after the second dose) (**Figure S2C**).

We went on to further evaluate the effects of WT and Omicron LNP-mRNA boosters in infectious virus neutralization assay, which closely correlated with pseudovirus neutralization results. The Omicron LNP-mRNA booster led to over 200-fold increase in neutralizing titers of infectious Omicron virus (**Figure 2D**), while WT booster induced a moderate increase (10-fold) in titers against Omicron live virus (**Figure 2D**). A significant boost of infectious Delta virus neutralizing titers was observed in mice receiving WT (12-fold) and Omicron (19-fold) LNP-mRNA boosters. A 20-fold difference in post-booster (day 180) neutralizing titers against infectious Omicron virus was observed between WT and Omicron booster groups (**Figure 2D**). Together, these data suggest that while both WT LNP-mRNA and Omicron LNP-mRNA boosters can strengthen the waning immunity; however, the heterologous booster with Omicron-specific mRNA vaccination (WT x2 + Omicron x1) has an effect significantly stronger than the homologous booster (WT x 3) against the live virus of Omicron variant, with comparable activity against the Delta variant.

Overall, the ELISA titers, pseudovirus and infectious virus neutralization activity were significantly correlated with each other across all groups and animals tested (**Figure S9**). These data suggested that a single dose of Omicron LNP-mRNA heterologous booster not only induced more potent anti-Omicron antibody response than WT booster, but also elicited broad activity against the WA-1 and Delta variant, in mouse models at the timepoints measured.

### Cross reactivity and epitope characterization of plasma antibodies from homologous Omicron mRNA, WT mRNA or heterologous WT + Omicron mRNA vaccination schemes

In light of the broad activity elicited by heterologous vaccination of WT and Omicron LNP-mRNA, we ask if these vaccination schemes can induce antibody responses against other SARS-CoV-2 variants and other pathogenic *Betacoronavirus* species. We sought to answer these questions by characterizing and comparing the anti-coronavirus cross reactivity conferred by Omicron mRNA vaccination alone, WT mRNA vaccination alone (homologous booster), or their uses in combination (Omicron mRNA vaccination as a heterologous booster on top of WT mRNA vaccination). The cross reactivity was evaluated using six spike RBDs, including SARS-CoV-2 WA-1, Beta (lineage B.1.351) variant, Delta variant, Omicron variant, SARS-CoV spike RBD (SARS RBD) and MERS-CoV spike RBD (MERS RBD). Two doses of Omicron LNP-mRNA induced high titers of antibodies that cross reacted with all spike RBDs tested except for MERS RBD, which shared low sequence identity (<40%) to SARS or SARS-CoV-2 spikes (**Figure 3A**). The antibody titer against SARS RBD was significantly lower than those against SARS-CoV-2 WA-1 or variants (**Figure S10A**). Among the SARS-CoV-2 variants characterized, the antibody response to Delta variant by Omicron LNP-mRNA was slightly weaker than others. Both WT and Omicron boosters after WT LNP-mRNA prime and boost led to potent antibody response to SARS-CoV and SARS-CoV-2 Beta variant (**Figure 3B**), while the response to MERS RBD was negligible and similar to PBS control. Within each ELISA antigen except for MERS RBD and Omicron RBD, the antibody response post WT or Omicron boosters (3 shots total) was numerically higher than that of plasma samples post a two-dose Omicron vaccine (Omicron x 2) (**Figure S10C**).

**Figure 3.**
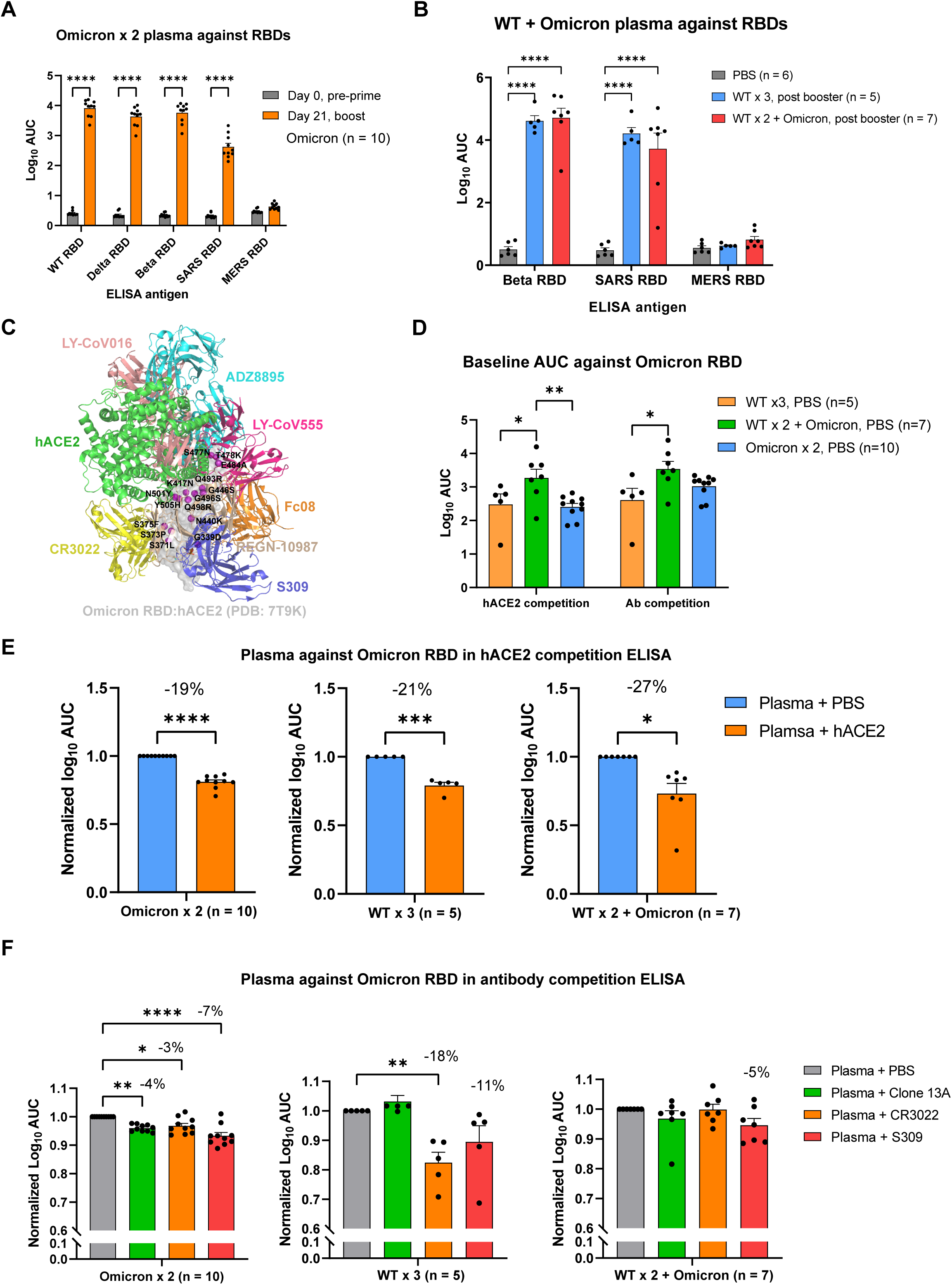
Cross reactivity and targeting sites characterization of plasma antibodies elicited by Omicron and WT LNP-mRNAs against SARS-CoV-2 VoCs and *Betacoronavirus* species. **A,** cross reactivity of plasma antibody from mice immunized with Omicron LNP mRNA (prime and boost) to SARS-CoV-2 VoCs and pathogenic coronavirus species (n = 10). **B**, cross reactivity of plasma antibody from mice immunized with WT (WT x 3) or Omicron (WT x 2 + Omicron) boosters to SARS-CoV-2 beta variant and pathogenic coronavirus species (n = 6 in PBS, n=5 in WT x 3, n = 7 in WT x 2 + Omicron). **C,** representative antibodies from major classes of RBD epitopes were shown by aligning spike RBDs in each of complex structures. The Omicron RBD surface was set to semi-transparent to visualize 15 RBD mutations and their relative positions to antibody epitopes. **D,** baseline titers of plasma from mice of different vaccination status (WT x 3, WT x 2 + Omicron, Omicron x 2) were shown as log_10_ AUC determined in hACE2 and antibody competition ELISA. Each group sample number is denoted with n (n = 10 in Omicron x 2, n=5 in WT x 3, n = 7 in WT x 2 + Omicron) in two independent assays (hACE2 and antibody competition ELISA). **E**, significant portion of plasma antibody from mice receiving Omicron (Omicron x 2, left panel) or WT + Omicron (WT x 3 middle, or WT x 2 + Omicron, right panel) LNP-mRNA competed with hACE2 for Omicron RBD binding in ELISA (n = 10 in Omicron x 2, n=5 in WT x 3, n = 7 in WT x 2 + Omicron). **F**, plasma antibody from mice receiving Omicron (Omicron x 2, n = 10, left panel) or WT + Omicron (WT x 3, n = 5, middle or WT x 2 + Omicron, n = 7, right panel) LNP-mRNA showed various extent of binding reduction in the presence of blocking antibodies with known epitopes on RBD. Data on dot-bar plots are shown as mean ± s.e.m. with individual data points in plots. Two-way ANOVA with Tukey’s multiple comparisons test and one-way ANOVA with Dunnett’s multiple comparisons were used to assess statistical significance. Statistical significance labels: * p < 0.05; ** p < 0.01; *** p < 0.001; **** p < 0.0001. Non-significant comparisons are not shown, unless otherwise noted as n.s., not significant.

A number of studies have shown that antibodies whose epitopes overlap with hACE2-binding motif were largely escaped by RBD mutations in variants of concerns, while antibodies whose epitopes fall outside the hACE2-binding motif were rarer and often exhibit broad neutralizing activity to SARS-like *Betacoroanviruses* (*Sarbecoviruses*)^35–38^. Because of such correlation between antibody epitope and cross reactivity, we performed competition ELISA using hACE2 or antibodies with known epitopes as competing agents to evaluate the epitopes, population and affinity of plasma antibodies elicited by Omicron or WT LNP-mRNA. The epitopes of RBD can be categorized into several major classes based on cluster analysis of available neutralizing antibody-RBD complex structures^38–42^. We displayed representative antibodies in each major epitope classes by aligning them with the recently solved Omicron RBD:hACE2 complex structure^31,32^ (**Figure 3C**). We then performed hACE2 and antibody competition ELISA using hACE2, Clone 13A, S309 and CR3022 as competing reagents to see if and to what extent group A-D^35^ (class I-III^36^, epitopes overlapped with hACE2) and group E-F (class IV, S309 and CR3022) antibodies were induced by these immunization schemes. Low-density Omicron RBD was coated in ELISA plate to ensure adequate competition between plasma antibodies and competing hACE2 or antibodies. In two independent experiments (hACE2 and antibody competition assays), the baseline titer of heterologous Omicron booster treated mice (WT x 2 + Omicron) in the absence of competing reagents was significantly higher than those of homologous WT booster treated mice (WT x 3), or mice receiving Omicron vaccination alone (Omicron x 2) (**Figure 3D**). Addition of high concentration hACE2 (Methods) resulted in a significant reduction of plasma antibody titers in mice vaccinated with Omicron (Omicron x2), WT (WT x 3) or WT + Omicron (WT x2 + Omicron) LNP-mRNA (**Figure 3E**). In the antibody competition assay, we used three antibodies with known RBD epitopes. Two of them (CR3022 and S309) are well-characterized representative antibodies from non-hACE2 competing classes. The Clone 13A is a humanized neutralizing antibody developed in our lab previously ^43^ and has an epitope that overlaps with the hACE2 binding motif. All three antibodies led to a significant decrease of plasma titers from Omicron vaccinated mice (Omicron x 2), while only CR3022 and S309 mediated a titer reduction in WT booster group (WT x 3) (**Figure 3F**). The WT + Omicron heterologous vaccination group showed minimal titer changes to all three antibodies (**Figure 3F**). These data suggested that a significant percentage of the pool of antibodies elicited by Omicron- or WT-vaccination shared binding epitopes with hACE2. In addition, antibody competition ELISA showed that both Omicron LNP-mRNA and WT LNP-mRNA vaccinated animals contained plasma antibodies targeting rare epitopes in class IV (or group E/F), which often exhibit broad activity against *Sarbecoviruses*.

## Discussion

The rapid spread of Omicron around the world, especially in countries with wide coverage of vaccines designed based on the ancestral antigen (e.g. WT mRNA vaccine), is particularly concerning. The extensive mutations in the Omicron spike gene mark a dramatic alteration in its antigenicity^15^. Omicron has high transmissibility and high level of immune evasion from WT mRNA vaccine induced immunity, which was reported from various emerging literature ^6,14–17^. Omicron’s strong association with reinfection^6^ or breakthrough infection^8,12^ and its heavily altered antigenicity prompted the idea of developing Omicron-specific mRNA vaccine.

To date (as of Feb 20, 2022), 4.35 billion people, i.e. 56% of the global population, received COVID-19 vaccination (Our World in Data^44^). Almost all those vaccines were designed based on the antigen from the ancestral virus, including the two approved mRNA vaccine BNT162b2 ^21^ and mRNA-1273 ^22^. Individuals receiving existing COVID-19 vaccines have waning immunity overtime^45–48^. Consistent with past reports, we observed a dramatic time-dependent decrease (around 40-fold) of antibody titers against Omicron, Delta variants and WA-1 strains 3 months after the second dose of WT mRNA vaccine in mice. This observed waning immunity is particularly concerning in the scenario of rapid spreading of Omicron variant, which largely escapes the humoral immune response elicited by WT mRNA vaccines as evident in published studies^14–16,18^ as well as in our current data. A recent report showed waning immunity in vaccinated individuals^24^ and that a booster shot using the WT based mRNA vaccine helps recover partial immunity. Our data showed that the neutralizing antibody titers after the boost with a WT based vaccine were still lower against Omicron than against WA-1 and other variants, urging for development and testing of an Omicron-specific vaccine. Vaccinee receiving heterologous vaccination of WT and Omicron LNP-mRNA have been exposed to both antigens and may have robust antibody response against cognate strains and other VoCs. Thus, it is important to evaluate and compare the immunogenicity of Omicron-specific vaccine candidate with WT vaccine as booster shots on top of two doses of WT mRNA vaccine. In fact, very recently, both Pfizer and Moderna have started their clinical trials to evaluate the efficacy of Omicron-specific mRNA vaccine in either homologous or heterologous vaccination settings^49–51^. Moderna has released an updated Phase 2/3 clinical trial for their Omicron-specific mRNA vaccine (mRNA-1273.529) along with the WT vaccine mRNA-1273 against COVID-19 Omicron variant (NCT05249829). The scale and swiftness of initiating these clinical trials exemplify the clinical importance and urgent need of curbing the Omicron surge and evaluating the Omicron-specific mRNA vaccine.

In this study, we generated a HexaPro-version full-length Omicron spike LNP-mRNA vaccine candidate. In mouse models, we found that it can induce potent Omicron-specific and broad anti-Sarbecovirus antibody response. With this vaccine candidate, we compared its boosting effect with WT counterpart on animals that previously received two-dose WT mRNA vaccine. An observation is that a single dose of WT or Omicron boosters significantly strengthened the waning immunity against Omicron and Delta variants. A number of recent preprints generated and tested Omicron-specific vaccine candidates, which had different vaccine antigen designs, compositions, and showed varying results of antibody responses alone or as boosters ^52–57^ (briefly summarized in **Table S1**). Three of them focused on evaluation of Omicron RBD mRNA vaccine alone in mice through neutralization assay and reported antibody response against Omicron but not other variants^52,55,56^. Two studies characterized the Omicron full-length spike mRNA stabilized by two proline mutations (S-2P) and compared their boosting efficacy with WT vaccine in mice^53^ and macaque^54^. Preprints Gagne et. al. and Ying et. al have shown that both WT and Omicron full-length spike mRNA boosters provided equivalent protection from Omicron challenge in non-human primates (NHPs) ^54^ or mice^53^. These results shared some commonalities, i.e. the effectiveness of an Omicron-specific vaccine; however, they diverged in the specific titers, as well as in the difference between WT- and Omicron-specific vaccines, potentially due to differences in vaccine antigen designs, compositions, modifications, experimental settings, animal models, or a combination of factors. Our study evaluated the potency of an Omicron-specific full-length spike mRNA vaccine with HexaPro mutations^27^, which were shown to stabilize the spike in prefusion state. Through well-correlated data from ELISA, pseudovirus and infection virus neutralization assays, we showed that both WT and Omicron boosters significantly restored waning immunity against Omicron and Delta variants. Interestingly, without sacrificing potency against Delta, heterologous Omicron booster achieved significantly higher neutralizing titers against Omicron than homologous WT booster. This observation is in line with findings from heterologous booster vaccination of different COVID-19 vaccines in clinical trials^25,26^. The broad anti-coronavirus activity after homologous or heterologous boosting was likely associated with plasma antibodies in rarer epitope classes, as observed in competition ELISA.

It has been shown that the neutralizing antibody level is highly predictive of immune protection from SARS-CoV-2 infection and the initial neutralization level is associated with decay of vaccine efficacy over time^58^. Compared to WT booster, we found that Omicron booster group consistently showed 10-20 fold higher titers against Omicron variant in ELISA, pseudovirus and infectious virus neutralization assays. Within the WT vaccinated group, the titer contrast against Omicron vs. Delta variants persisted over time. Omicron-booster group have been exposed to both WT and Omicron antigens and showed equally potent titers against Omicron and Delta. While our study is in animals, the antibody responses to vaccination are conserved between mouse and human, highlighted by the fact that mice are the main preclinical model used by vaccine developers ^59,60^.

The titer against Omicron by single dose Omicron LNP-mRNA was similar to that observed 2 weeks post boost of WT LNP-mRNA (log10 AUC or log10 IC50 around 3), although it is still unclear whether the potency of the Omicron mRNA vaccine is associated with the high number of Omicron mutations. As various extent of cross reactivity was observed among WT and/or Omicron vaccinated animals, we sought to understand their cross-reactive immunity by characterizing vaccine-elicited antibody epitopes and population through competition ELISA. In the Omicron RBD competition ELISA, the baseline titer of Omicron LNP-mRNA booster group (WT x2 + Omicron) was significantly higher than WT booster (WT x 3) or Omicron LNP-mRNA (Omicron x 2), which may explain its lower susceptibility to the block of competing antibodies. All three vaccination groups showed significant titer reduction in presence of hACE2, suggestive of abundant plasma antibody population sharing hACE2 binding epitopes, which are often associated with immune escape by variants mutations. The plasma from mice vaccinated with two doses of Omicron LNP-mRNA (Omicron x 2) or three doses of WT LNP-mRNA (WT x 3) exhibited comparable baseline titers and significant titer decrease when co-incubated with CR3022 or S309 blocking antibodies, indicating the existence of plasma antibody population sharing group E/F^35^ or class IV^36^ epitopes. Because of their similar baseline titers, the greater titer reduction in WT booster group may stem from larger population of group E/F antibodies, which was associated higher cross-reactive response against SARS RBD (Figure S10C). Albeit insignificant, the titer change of Omicron booster group (WT x 2 + Omicron) by S309 antibody was greatest among three competing antibodies, hinting a role of epitope IV antibodies in the cross immunity elicited by heterologous vaccination of WT and Omicron LNP-mRNA.

In summary, this study generated an Omicron-specific HexaPro spike LNP-mRNA vaccine candidate, studied its immunogenicity, and compared it with the WT counterpart in the context of previously WT vaccinated animals. Our results showed that a single dose of either a homologous booster with WT LNP-mRNA or a heterologous booster with Omicron LNP-mRNA restored the waning antibody response, with over 200-fold titer increase by Omicron boosters. Interestingly, the heterologous Omicron LNP-mRNA booster elicited Omicron neutralizing titers higher than the homologous WT booster. The heterologous Omicron booster shot provided strong neutralizing antibody response against Omicron variant and comparable humoral antibody against WA-1 and Delta variants. All three types of vaccination, including Omicron mRNA alone, WT mRNA alone, and Omicron as a heterologous booster on top of WT mRNA, elicited broad antibody responses, including activities against SARS-CoV-2 VoCs, as well as other *Betacoronavirus* species such as SARS-CoV, but not MERS-CoV. Together, these data provided direct proof-of-concept assessments of Omicron-specific mRNA vaccination *in vivo*, both alone and as a heterologous booster to the existing widely-used mRNA vaccine form.

## Acknowledgments

We thank various members from our labs for discussions and support. We thank staffs from various Yale core facilities (Keck, YCGA, HPC, YARC, CBDS and others) for technical support. We thank Drs. Tsemperouli, Karatekin, and others for providing equipment and related support. We thank Drs. Lucas and Iwasaki for sharing the Omicron virus isolate. We thank various support from Department of Genetics; Institutes of Systems Biology and Cancer Biology; Dean’s Office of Yale School of Medicine and the Office of Vice Provost for Research.

## Funding

This work is supported by DoD PRMRP IIAR (W81XWH-21-1-0019) and discretionary funds to SC; and Ludwig Foundation, Mathers Foundation, Burroughs Wellcome Fund to CBW. The TEM core is supported by NIH grant GM132114 to CL.

## Institutional Approval

This study has received institutional regulatory approval. All recombinant DNA (rDNA) and biosafety work were performed under the guidelines of Yale Environment, Health and Safety (EHS) Committee with approved protocols (Chen 18-45, 20-18, 20-26). All animal work was performed under the guidelines of Yale University Institutional Animal Care and Use Committee (IACUC) with approved protocols (Chen-2020-20358; Chen 2021-20068; Wilen 2021-20198).

## Methods

### Molecular cloning

The Omicron spike amino acid sequence was derived from two lineage BA.1 Omicron cases identified in Canada on Nov.23^rd^, 2021 (GISAID EpiCoV, EPI_ISL_6826713 and EPI_ISL_6826714). Omicron spike cDNA were codon optimized, synthesized as gblocks (IDT) and cloned to mRNA vector with 5’, 3’ untranslated region (UTR) and poly A tail. The furin cleave site (RRAR) was replaced with a GSAS short stretch in the mRNA vector. HexaPro mutations were introduced in the WT sequence (Wuhan-Hu-1, which was used for the current clinical mRNA vaccines) and Omicron variant spike sequence of mRNA vector to improve expression and prefusion state^27^. The accessory plasmids for pseudovirus assay including pHIVNLGagPol and pCCNanoLuc2AEGFP were from Dr. Bieniasz’ lab^61^. The C-terminal 19 amino acids were deleted in the SARS-CoV-2 spike sequence for the pseudovirus assay.

### Cell Culture

HEK293T, HEK293FT (Thermo Fisher) and 293T-hACE2 (gifted from Dr Bieniasz’ lab) cell lines were maintained in Dulbecco’s modified Eagle’s medium (DMEM, Thermo fisher) supplemented with 10% Fetal bovine serum (Hyclone) and 1% penicillin-streptomycin (Gibco, final concentration penicillin 100 unit/ml, streptomycin 100 µg/ml), which is denoted as complete growth medium. Cells were split every 2 days at a split ratio of 1:4 when the confluency reached over 80%. Vero-E6 cells were cultured in Dulbecco’s Modified Eagle Medium (DMEM) with 5% heat-inactivated fetal bovine serum (FBS).

### In vitro mRNA transcription and vaccine formulation

A Hiscribe^TM^ T7 ARCA mRNA Kit (with tailing) (NEB, Cat # E2060S) was used to in vitro transcribe codon-optimized mRNA encoding HexaPro spikes of SARS-CoV-2 WT and Omicron variant with 50% replacement of uridine by N1-methyl-pseudouridine. The DNA template was linearized before mRNA transcription and contained 5′ UTR, 3′ UTR and 3’polyA tail as flanking sequence of spike open reading frame.

The purified mRNA was generated by following NEB manufacturer’s instructions and kept frozen at -80 °C until further use. The lipid nanoparticles mRNA was assembled using the NanoAssemblr® Ignite™ instrument (Precision Nanosystems) according to manufacturers’ guidance. In brief, lipid mixture was mixed with prepared mRNA in 25mM sodium acetate at pH 5.2 on Ignite instrument at a molar ratio of 6:1 (LNP: mRNA), similar to previously described^59,62^. The LNP encapsulated mRNA (LNP-mRNA) was buffer exchanged to PBS using 100kDa Amicon filter (Macrosep Centrifugal Devices 100K, 89131-992). Sucrose was added as a cryoprotectant. The particle size of mRNA-LNP was determined by DLS device (DynaPro NanoStar, Wyatt, WDPN-06) and TEM described below. The encapsulation rate and mRNA concentration were quantified by Quant-iT™ RiboGreen™ RNA Assay (Thermo Fisher).

### Validation of LNP-mRNA mediated spike expression *in vitro* and receptor binding capability of expressed Omicron HexaPro spikes

On day 1, HEK293T cells were seeded at 50% confluence in 24-well plate and mixed with 2 µg Omicron LNP-mRNA. After 16 hours, the cells were collected for flow cytometry. The spike expression on cell surface were detected by staining cells with human ACE2–Fc chimera (Sino Biological, 10108-H02HG) in MACS buffer (D-PBS with 2 mM EDTA and 0.5% BSA) for 20 min on ice. Cells were washed twice after the primary stain and incubated with PE–anti-human Fc antibody (Biolegend, 410708) in MACS buffer for 20 min on ice. During secondary antibody staining, live/Dead aqua fixable stain (Invitrogen) was used to assess cell viability. Data was collected on BD FACSAria II Cell Sorter (BD) and analyzed using FlowJo software.

### Negative-stain TEM

Formvar/carbon-coated copper grid (Electron Microscopy Sciences, catalog number FCF400-Cu-50) was glow-discharged and covered with 6 µl of the sample for 1 min before blotting away the sample. The sample was double-stained with 6 µl of 2% (w/v) uranyl formate (Electron Microscopy Sciences, catalog number 22450) for 5 seconds (first stain) and 1 min (second stain), blotting away after each stain. Images were collected using a JEOL JEM-1400 Plus microscope with an acceleration voltage of 80 kV and a bottom-mount charge-coupled device camera (4k by 3k, Advanced Microscopy Technologies).

### Mouse vaccination

Vaccine immunogenicity study used 6 weeks old female C57BL/6Ncr (B6) mice purchased from Charles River. The mice-housing condition was maintained at regular ambient room temperature (65-75°F, or 18-23°C), 40-60% humidity, and a 14 h:10 h day/night cycle. Each mice cage was individually ventilated with clean food, water, and bedding. Two sets of immunization experiments were performed: vaccination with Omicron LNP-mRNA, and sequential vaccination with WT LNP-mRNA, followed by WT or Omicron LNP mRNA booster. For the Omicron LNP-mRNA vaccination experiment, five mice were immunized with 10 µg Omicron LNP-mRNA on day 0 (prime) and day 14 (boost). Retro-orbital blood was collected prior to vaccine injection on day 0, day 13 and day 21. For WT and Omicron LNP-mRNA sequential vaccination experiment, three mice were administered with either 100 µl PBS or two-dose 1 µg WT (21 days apart) and 10 µg Omicron LNP-mRNA (14 days apart). Retro-orbital blood was collected prior to vaccine injection on day 35, day 127, day 140 and day 148.

### Isolation of plasma and PBMCs from blood

At the defined time points, retro-orbital blood was collected from mice. The isolation of PBMCs and plasma was achieved via centrifugation using SepMate-15 and Lymphoprep gradient medium (StemCell Technologies). 200 µl blood was immediately diluted with 800 µl PBS with 2% FBS. The blood diluent was then added to SepMate-15 tubes with 6ml Lymphoprep (StemCell Technologies). Centrifugation at 1200 g for 20 minutes was used to isolate RBCs, PBMCs and plasma. 250ul diluted plasma was collected from the surface layer. The remaining solution at the top layer was poured to a new tube to isolate PBMCs, which were washed once with PBS + 2% FBS. The separated plasma was used in ELISA and neutralization assay.

### ELISA

3 µg/ml of spike antigens were coated onto the 384-well ELISA plates (VWR, Cat # 82051-300) overnight at 4 degree. The antigen panel used in the ELISA includes RBDs of SARS RBD (AcroBiosystems, SPD-S52H6), MERS RBD (AcroBiosystems, SPD-M52H6), 2019-nCoV WA-1 (Sino Biological 40592-V08B), Delta variant B.1.617.2 (Sino Biological 40592-V08H90), Beta variant B.1.351 (Sino Biological 40592-V08H85) and Omicron variant B.1.1.529 (Sino Biological 40592-V08H121). Plates were washed with PBST (PBS plus 0.5% Tween 20) three times in the 50TS microplate washer (Fisher Scientific, NC0611021) and blocked with 0.5% BSA in PBST at room temperature for one hour. Plasma was fourfold serially diluted starting at a 1:500 dilution. Diluted plasma samples were added to the plates and incubated at room temperature for one hour, followed by washes with PBST five times. Anti-mouse secondary antibody (Fisher, Cat# A-10677) at 1:2500 dilution in blocking buffer was incubated at room temperature for one hour. Plates were washed five times and developed with tetramethylbenzidine substrate (Biolegend, 421101). The reaction was stopped with 1 M phosphoric acid after 20 min at room temperature, and OD at 450 nm was measured by multimode microplate reader (PerkinElmer EnVision 2105). The binding response (OD450) was plotted against the dilution factor in log10 scale as the dilution-dependent response curve. The area under curve of the dilution-dependent response (Log10 AUC) was calculated to quantify the potency of the plasma antibody binding to spike antigens. The fold change of antibody titer was estimated using this equation: ratio = 10 ^ (AUC1 - AUC2)

### hACE2 and antibody competition ELISA

The 384-well plate was coated with 0.6 µg/ml Omicron RBD at 4 degree overnight before washed with PBST (0.5% Tween-20) three times and blocked with 2% BSA in PBST for 1 hour at room temperature. In hACE2 and antibody competition ELISA, 15 µg/ml hACE2 (Sino, 10108-H08H) or 10 µg/ml antibodies including Clone 13A (Chen lab), CR3022 (Abcam, Ab273073) and S309 (BioVision, A2266) were respectively added to the plate 1 hour prior to subsequent incubation with serially diluted plasma for another hour at room temperature. After coincubation of plasma and hACE2/antibodies, the plate was washed five times with PBST and incubated with anti-mouse secondary antibody with minimal cross reactivity with human IgG (Biolegend, 405306). The plate was washed five times after 1-hour secondary antibody incubation and developed with tetramethylbenzidine substrate (Biolegend, 421101). The reaction was stopped with 1 M phosphoric acid after 20 min at room temperature, and OD at 450 nm was measured by multimode microplate reader (PerkinElmer EnVision 2105). The normalized AUC was calculated by normalizing the value with AUC determined in PBS group.

### Omicron, WA-1 and Delta pseudovirus production and characterization

For the neutralization assay, HIV-1 based SARS-CoV-2 WA-1, B.1.617.2 (Delta) variant, and B.1.1.529 (Omicron) variant pseudotyped virions were packaged using a coronavirus spike plasmid, a reporter vector and a HIV-1 structural protein expression plasmid. The reporter vector, pCCNanoLuc2AEGFP, and plasmid expressing HIV-1 structural proteins (pHIVNLGagPol) were gifts from Dr Bieniasz’s lab. The spike plasmid for SARS-CoV-2 WA-1 pseudovirus truncated C-terminal 19 amino acids (denoted as SARS-CoV-2-Δ19) and was from Dr Bieniasz’ lab. Spike plasmids expressing C-terminally truncated SARS-CoV-2 B.1.617.2 variant S protein (Delta variant-Δ19) and SARS-CoV-2 B.1.1.529 variant S protein (Omicron variant-Δ19) were made based on the pSARS-CoV-2-Δ19. All pseudoviruses were produced under the same conditions. Briefly, 293FT cells were seeded in 150 mm plates, and transfected with 21 µg pHIVNLGagPol, 21 µg pCCNanoLuc2AEGFP, and 7.5 µg of corresponding plasmids, in the presence of 198 µl PEI (1mg/ml, PEI MAX, Polyscience). At 48 h after transfection, the supernatant was filtered through a 0.45-µm filter, and frozen in -80°C.

To characterize the titer of WA-1, Delta, and Omicron pseudoviruses packaged, 1 x10^4^ 293T-hACE2 cells were plated in each well of a 96-well plate. In the next day, different volumes of pseudovirus supplemented with culture medium to a total volue of 100 µL were added into 96-well plates with 293T-hACE2. Plates were incubated at 37°C for 24 hr. Then cells were washed with MACS buffer once and the percent of GFP-positive cells were counted by Attune NxT Acoustic Focusing Cytometer (Thermo Fisher). To normalize pseudovirus titer, 1 x10^4^ 293T-hACE2 cells were plated in each well of a 96-well plate. In the next day, 50 µL pseudovirus was mixed with 50 µL culture medium to 100 µL. The mixture was incubated for 1 hr in the 37 °C incubator, supplied with 5% CO2, and added into 96-well plates with 293T-hACE2. Plates were incubated at 37°C for 24 hr. Then cells were washed with MACS buffer once and the percent of GFP-positive cells were counted by Attune NxT Acoustic Focusing Cytometer (Thermo Fisher). Delta pseudovirus and Omicron pseudovirus were diluted accordingly to match the functional titer of WA-1 pseudovirus for neutralization assay of plasma samples.

### Pseudovirus neutralization assay

The SARS-CoV-2 pseudovirus assays were performed on 293T-hACE2 cells. One day before infection, 1 x10^4^ 293T-hACE2 cells were plated in each well of a 96-well plate. In the next day, plasma collected from mice were serially diluted by 5 fold with complete growth medium at a starting dilution of 1:100. 55 µL diluted plasma was mixed with the same volume of SARS-CoV-2 WA-1, Delta variant, or Omicron variant pseudovirus and was incubated for 1 hr in the 37 °C incubator, supplied with 5% CO2. 100 µL of mixtures were added into 96-well plates with 293T-hACE2. Plates were incubated at 37°C for 24 hr. Then cells were washed with MACS buffer once and the percent of GFP-positive cells were counted by Attune NxT Acoustic Focusing Cytometer (Thermo Fisher). The 50% inhibitory concentration (IC50) was calculated with a four-parameter logistic regression using GraphPad Prism (GraphPad Software Inc.). If the fitting value of IC50 is negative (i.e. negative titer), which suggested undetectable neutralization activity, the value was set to baseline (1, 0 in log scale).

### Omicron and Delta live virus production and characterization

Full-length SARS-CoV-2 Omicron (BA.1) and Delta (B.1.617.2) isolates were a gift of Carolina Lucas and Akiko Iwasaki, and were isolated as previously described^63^. To expand viral stocks, 107 Vero-E6 cells stably overexpressing ACE2 and TMPRSS2 were infected with SARS-CoV-2 at an MOI of approximately 0.01. The Omicron stock was collected 2 dpi, clarified by centrifugation (450 xg for 10 minutes), filtered through a 0.45-micron filter, and concentrated ten-fold using Amicon Ultra-15 columns. To increase titer, the Delta stock was collected at 1 dpi, clarified, filtered, and used to infect 5 x 107 Vero-E6 cells overexpressing ACE2 and TMPRSS2. At 1 dpi, supernatant was harvested, clarified, filtered and concentrated as above. Viral stocks were titered by plaque assay in Vero-E6 cells as previously described^64^.

### Infectious virus neutralization assay

The complements and other potential neutralizing agents were heat inactivated in mouse plasma prior to infectious virus neutralization assay. Mouse plasma samples were serially diluted, then incubated with SARS-CoV-2 Omicron live virus for 1 h at 37°C. The Omicron live virus was isolated from nasopharyngeal specimens and sequenced as part of the Yale SARS-CoV-2 Genomic Surveillance Initiative’s weekly surveillance Program in Connecticut^65^. After coincubation, plasma/virus mixture was added to Vero-E6 cells overexpressing ACE2/TMPRSS2. Cell viability was measured at 3dpi or 5dpi using CellTiter Glo.

### Standard statistics

Standard statistical methods were applied to non-high-throughput experimental data. The statistical methods are described in figure legends and/or supplementary Excel tables. The statistical significance was labeled as follows: n.s., not significant; * p < 0.05; ** p < 0.01; *** p < 0.001; **** p < 0.0001. Prism (GraphPad Software) and RStudio were used for these analyses. Additional information can be found in the supplemental excel tables.

### Schematic illustrations

Schematic illustrations were created with Affinity Designer or BioRender.

### Replication, randomization, blinding and reagent validations

Biological or technical replicate samples were randomized where appropriate. In animal experiments, mice were randomized by littermates.

Experiments were not blinded.

Commercial antibodies were validated by the vendors, and re-validated in house as appropriate.

Custom antibodies were validated by specific antibody - antigen interaction assays, such as ELISA.

Isotype controls were used for antibody validations.

Cell lines were authenticated by original vendors, and re-validated in lab as appropriate.

All cell lines tested negative for mycoplasma.

## Data availability

All data generated or analyzed during this study are included in this article and its supplementary information files. Specifically, source data and statistics are provided in a supplementary table excel file. No custom code was used in this study. Additional information related to this study are available from the corresponding author(s) upon reasonable request.

## Extended Data Figures

**Extended Data Figure 1 (Figure S1).**
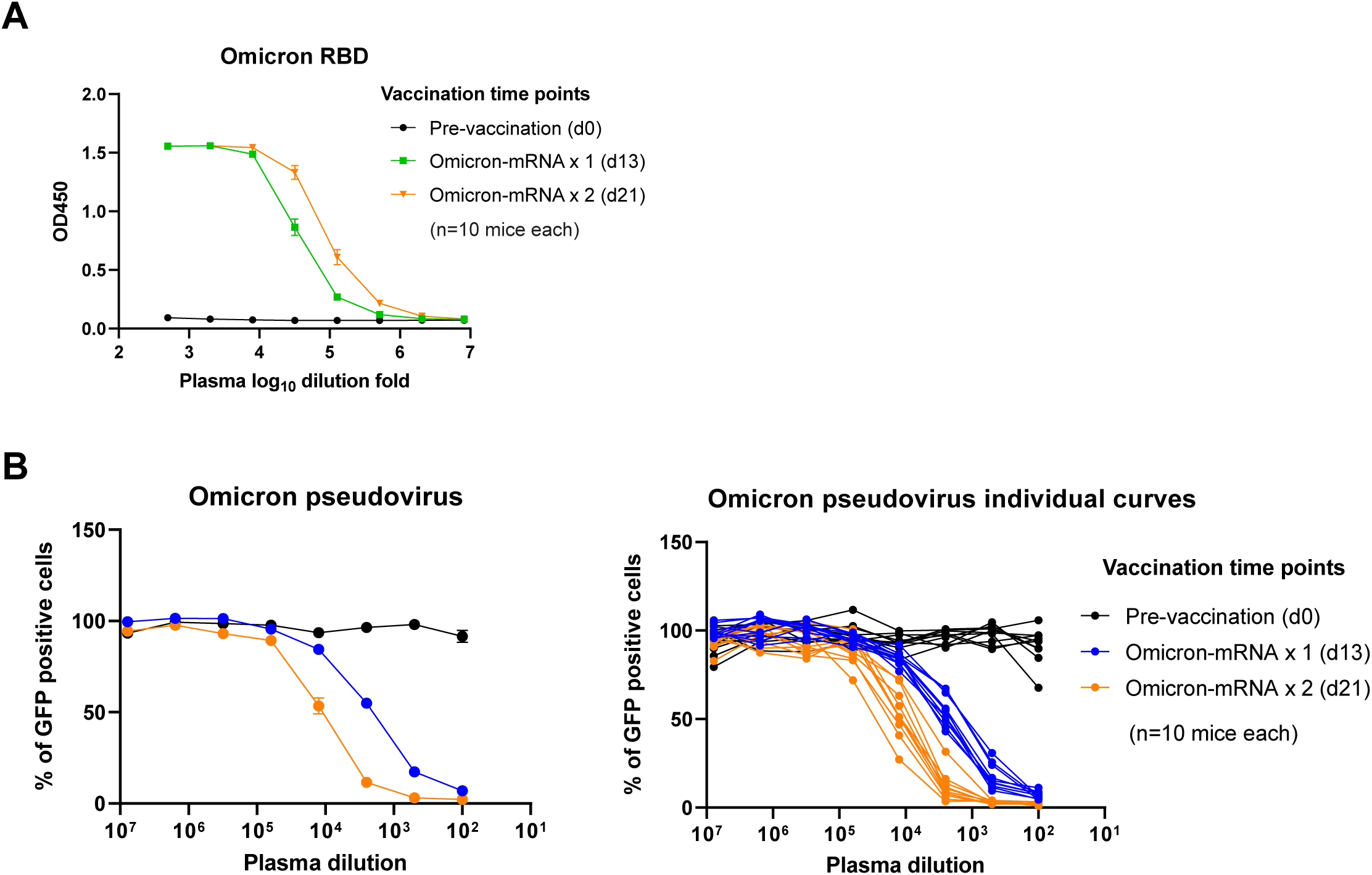
ELISA and neutralization titration curves over serial dilution of plasma collected at different timepoints from mice administered with PBS or WT and/or Omicron LNP-mRNA. **A,** ELISA titration curves over serial log_10_-transformed dilution points of plasma collected from mice before and after immunization with Omicron LNP-mRNA at defined time points (n = 10). Average curves, data are shown as mean ± s.e.m.. **B,** Omicron pseudovirus titration curves over serial log_10_-transformed dilution points of plasma collected from mice before and after immunization with Omicron LNP-mRNA at defined time points (n = 10). Left panel, average curves, data are shown as mean ± s.e.m.; Right panel, individual curves.

**Extended Data Figure 2 (Figure S2).**
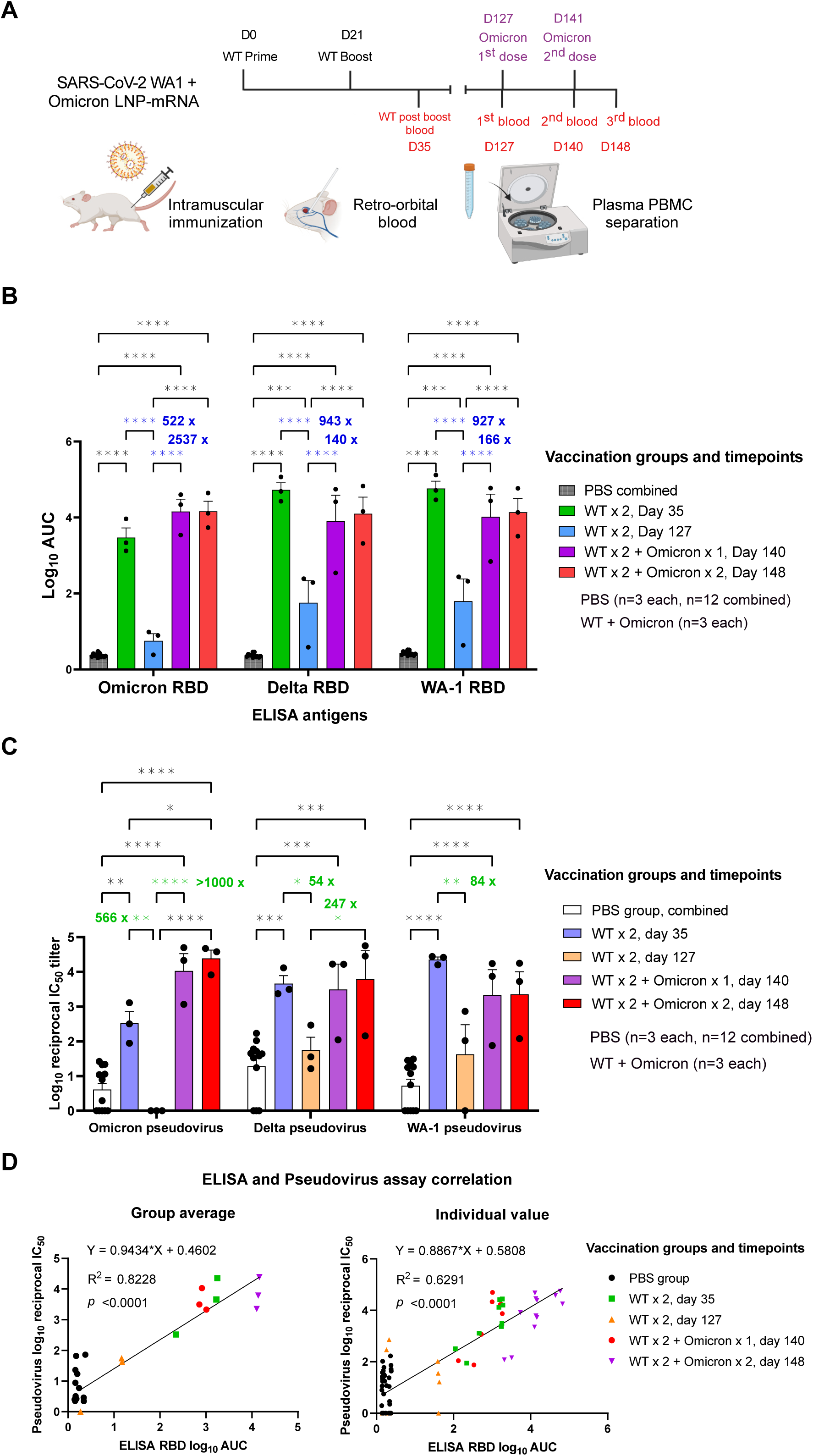
Both WT and Omicron specific LNP-mRNA booster shots greatly improved waning immunity of mice vaccinated with SARS-CoV-2 WT LNP-mRNA against SARS-CoV-2 Delta and Omicron variants (Independent experiment 1 or batch 1). **A**, Schematics showing the immunization and blood sampling schedule of mice administered with 1 µg WT LNP-mRNA prime (WT x 1) and boost (WT x 2) as well as 10 µg WT or Omicron-specific LNP-mRNA booster shots. The plasma and PBMCs were separate from blood for downstream assays. **B**, Bar graph comparing binding antibody titers of mice administered with PBS or WT and Omicron LNP-mRNA against Omicron, Delta and WT RBD (ELISA antigens). The antibody titers were quantified as Log_10_ AUC based on titrations curves in Extended Data Figure 1A. PBS sub-groups (n=3 each) collected from different matched time points showed no statistical differences between each other, and were combined as one group (n=9). **C**, Neutralizing antibody titers in the form of log_10_-transformed reciprocal IC50 calculated from fitting the titration curve with a logistic regression model (n = 3). **D**, Correlation of neutralization titers (log_10_ reciprocal IC50, y axis) and ELISA titers (log_10_ AUC, x axis) from matched vaccination group (left panel) or individual mouse (right panel). PBS samples from different timepoints were shown as one group in correlation map and were not included in linear regression model. Each dot in bar graphs represents value from one group average (left panel), or one individual mouse (right panel). Titer ratios were indicated in each graph and fold change is calculated from (ratio - 1). Data on dot-bar plots are shown as mean ± s.e.m. with individual data points in plots. Two-way ANOVA with Tukey’s multiple comparisons test was used to assess statistical significance. Statistical significance labels: * p < 0.05; ** p < 0.01; *** p < 0.001; **** p < 0.0001. Non-significant comparisons are not shown, unless otherwise noted as n.s., not significant.

**Extended Data Figure 3 (Figure S3).**
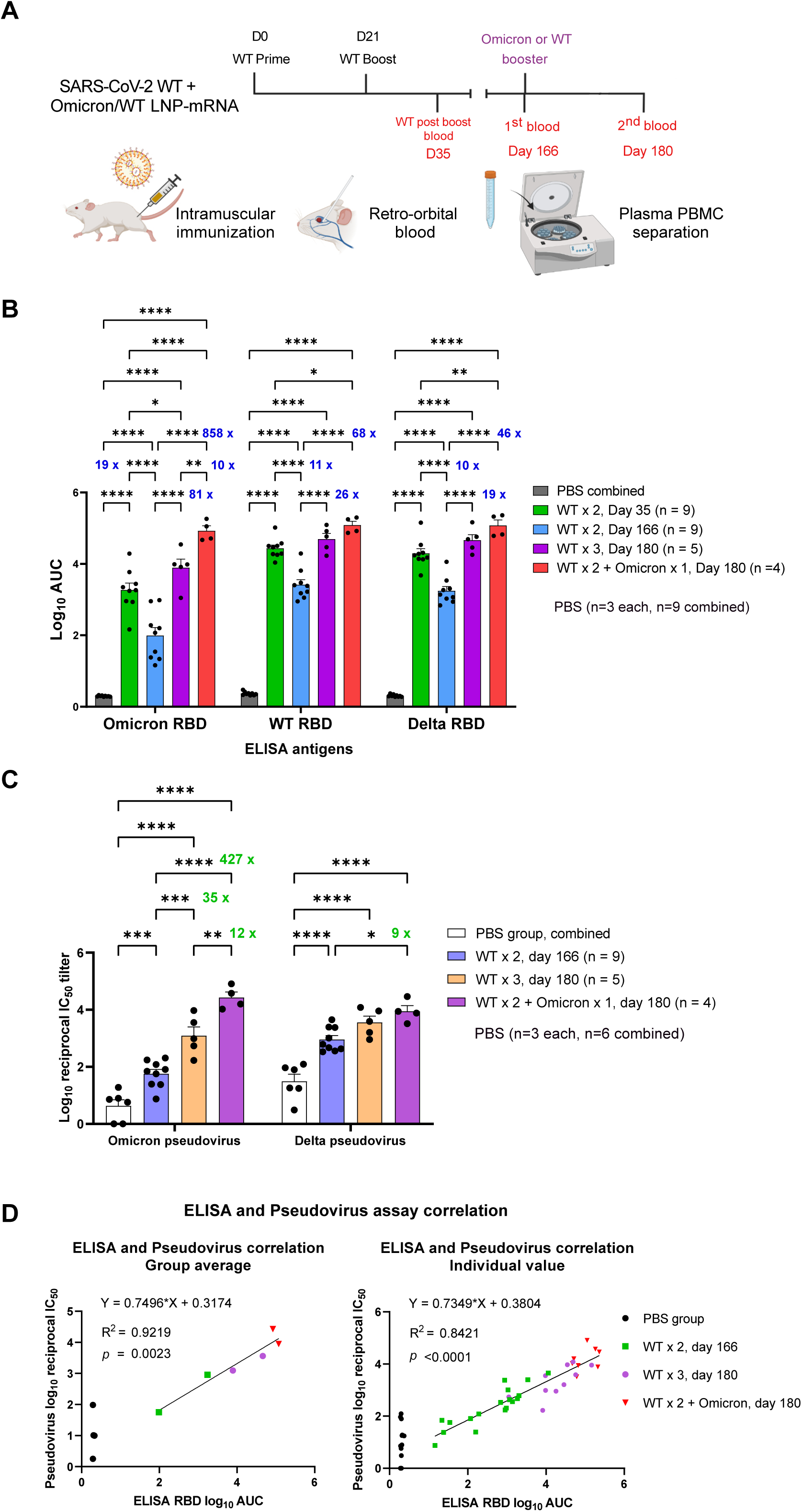
Omicron specific LNP-mRNA booster shots greatly improved waning immunity of mice vaccinated with SARS-CoV-2 WT LNP-mRNA against SARS-CoV-2 Delta and Omicron variants (Independent experiment 2 or batch 2). **A**, Schematics showing the immunization and blood sampling schedule of mice administered with 1 µg WT LNP-mRNA prime (WT x 1) and boost (WT x 2) as well as 10 µg Omicron-specific LNP-mRNA booster shots. The plasma and PBMCs were separate from blood for downstream assays. **B**, Bar graph comparing binding antibody titers of mice administered with PBS or WT and Omicron LNP-mRNA against Omicron, Delta and WT RBD (ELISA antigens). The antibody titers were quantified as Log_10_ AUC based on titrations curves in Extended Data Figure 1A. PBS sub-groups (n=3 each) collected from different matched time points showed no statistical differences between each other, and were combined as one group (n=6). **C**, Neutralizing antibody titers in the form of log_10_-transformed reciprocal IC50 calculated from fitting the titration curve with a logistic regression model (n = 9 before booster, n=5 in WT x 3, n = 4 in WT x 2 + Omicron). **D**, Correlation of neutralization titers (log_10_ reciprocal IC50, y axis) and ELISA titers (log_10_ AUC, x axis) from matched vaccination group (left panel) or individual mouse (right panel). PBS samples from different timepoints were shown as one group in correlation map and were not included in linear regression model. Each dot in bar graphs represents value from one group average (left panel), or one individual mouse (right panel). Titer ratios were indicated in each graph and fold change is calculated from (ratio - 1). Data on dot-bar plots are shown as mean ± s.e.m. with individual data points in plots. Two-way ANOVA with Tukey’s multiple comparisons test was used to assess statistical significance. Statistical significance labels: * p < 0.05; ** p < 0.01; *** p < 0.001; **** p < 0.0001. Non-significant comparisons are not shown, unless otherwise noted as n.s., not significant.

**Extended Data Figure 4 (Figure S4).**
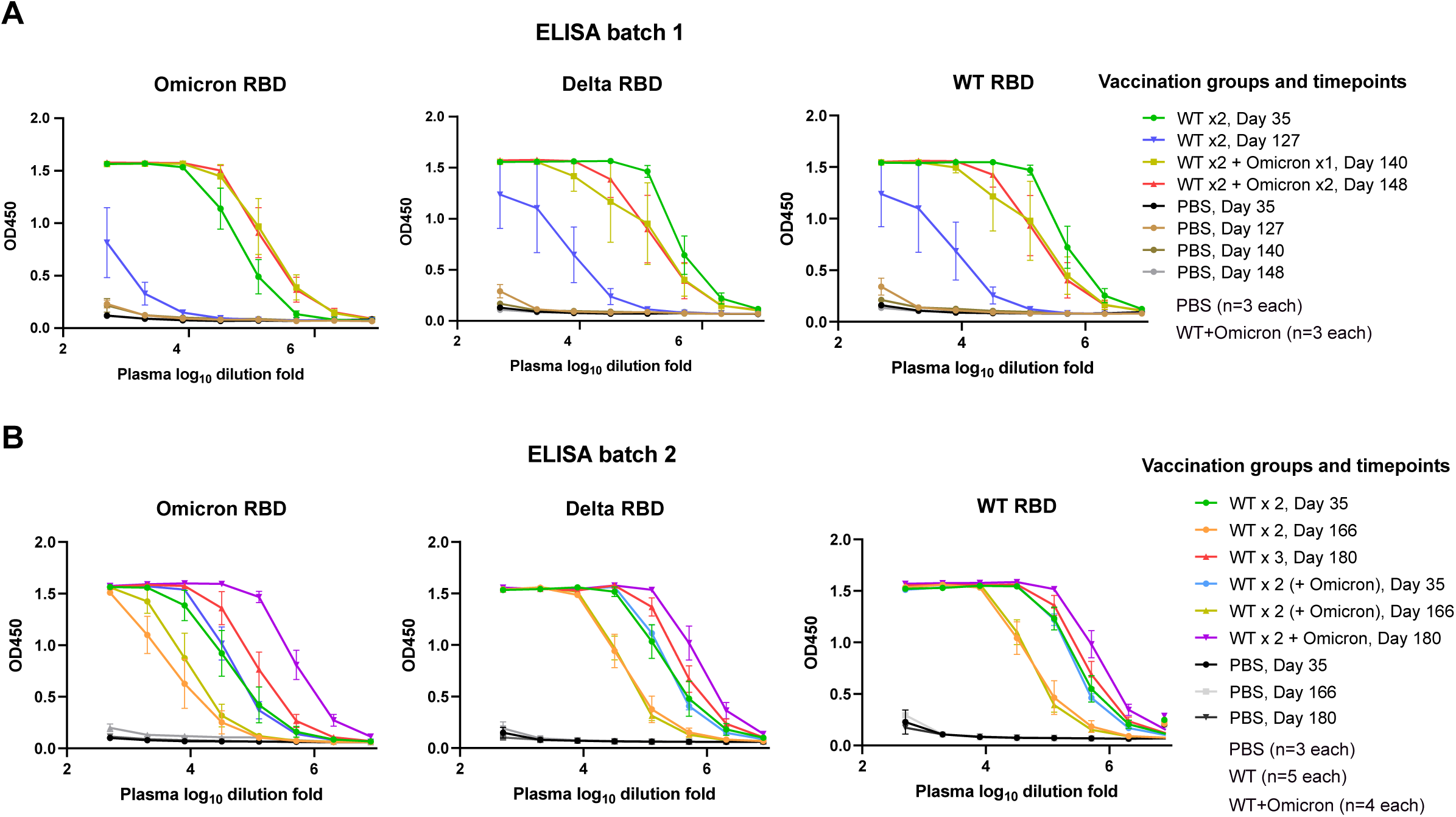
ELISA titration curves over serial dilution of plasma collected at different timepoints from mice administered with PBS or WT and/or Omicron LNP-mRNA. **A,** ELISA titration curves of batch 1 experiment (n = 3). **B**. ELISA titration curves of batch 2 experiment (n = 9 before booster, n=5 in WT x 3, n = 4 in WT x 2 + Omicron). The OD450 values were plotted against a series of log_10_-transformed dilution points of plasma from mice 35 days post WT prime, >4 months post WT prime (day 127 in batch 1 and day 166 in batch 2) and 2 weeks post booster (day 140 in batch 1 and day 180 in batch 2) of WT or Omicron LNP-mRNA, against spike receptor binding domain (RBD) antigens of Omicron variant (left), Delta (mid) and WT (right) were shown.

**Extended Data Figure 5 (Figure S5).**
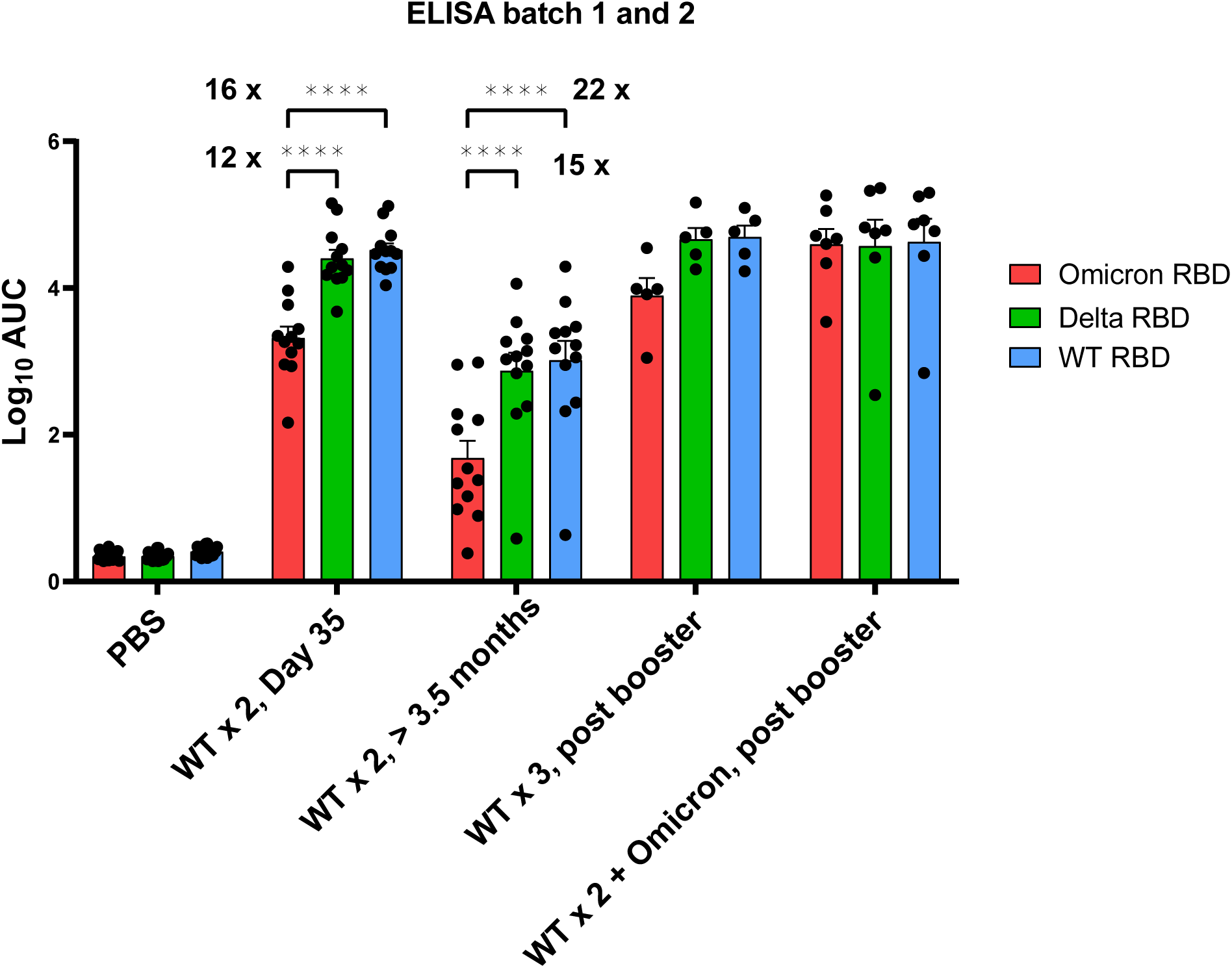
Binding antibody titers of mice administered with PBS or WT and Omicron LNP-mRNA against Omicron, Delta and WT RBD (ELISA antigens), were grouped by vaccination timepoints to compare titers against different RBD antigens. The antibody titers were quantified as area under curve of log_10_-transformed titration curve (Log_10_ AUC) in Extended Data Figure 2. The data were derived from independent experiment 1 and 2.

**Extended Data Figure 6 (Figure S6).**
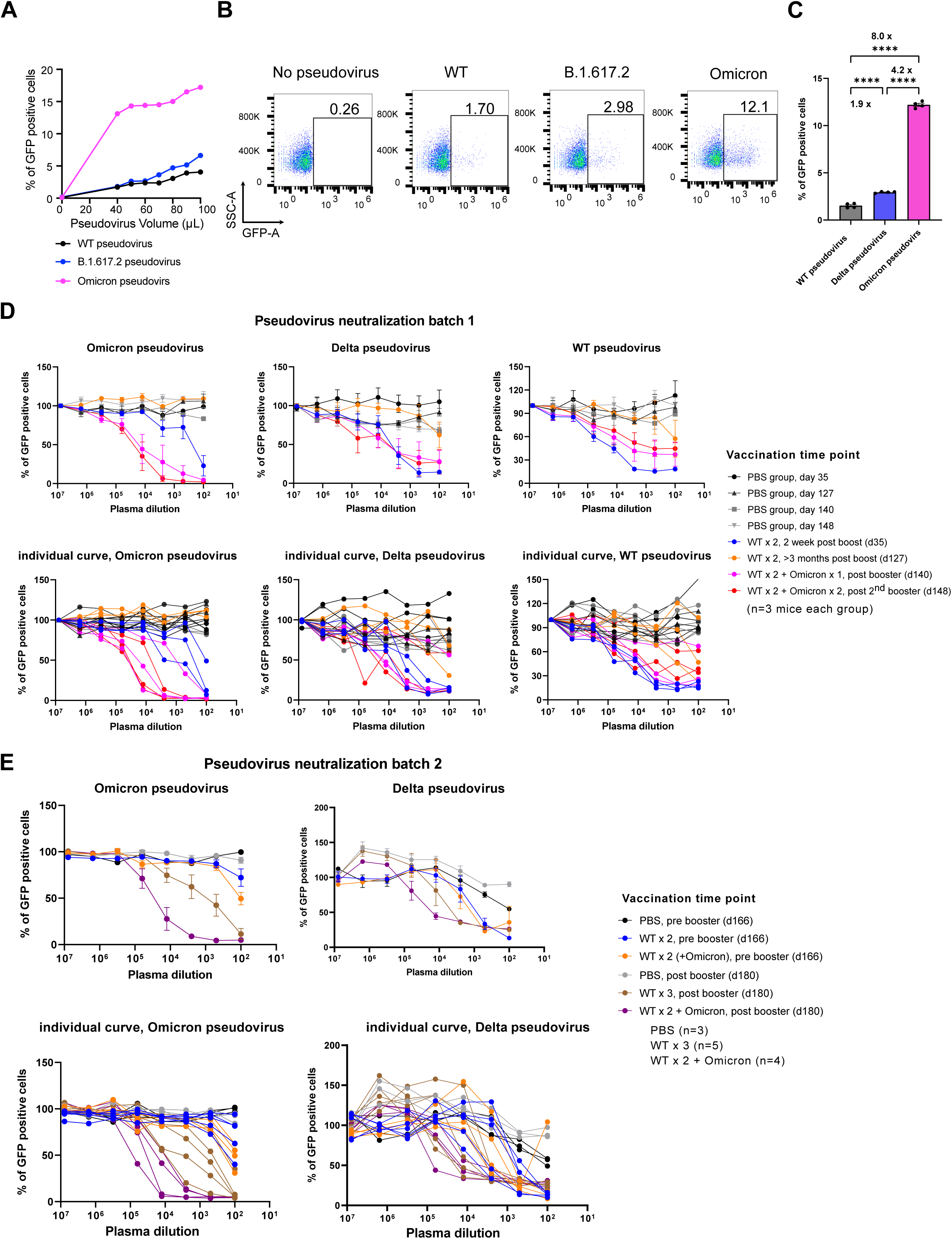
Omicron, Delta and WT pseudovirus production, characterization and neutralization assay. **A.** Functional titration curves of Omicron, Delta and WA-1 pseudoviruses in hACE2+ cells. **B.** Representative Flow Cytometry plots of infectivity of Omicron, Delta and WA-1 pseudoviruses in hACE2+ cells. **C.** Quantification of infectivity of Omicron, Delta and WT pseudoviruses in hACE2+ cells. **D,** Neutralization titration curves from batch 1 experiment (n = 3). **E.** Neutralization titration curves from batch 2 experiment (n = 9 before booster, n=5 in WT x 3, n = 4 in WT x 2 + Omicron). Percent of pseudovirus infected cells was plotted over serial dilutions of plasma from mice 35 days post WT prime, >4 months post WT prime (day 127 in batch 1 and day 166 in batch 2) and 2 weeks post booster (day 140 in batch 1 and day 180 in batch 2) of WT and Omicron LNP-mRNA against Omicron (left), Delta (mid) and WT (right) pseudovirus. Pseudovirus infection rate was calculated from percent of GFP positive cells and was plotted against plasma dilution (log_10_ transformed) as titration curve. Top panels, average curves, data are shown as mean ± s.e.m.; Bottom panels, individual curves.

**Extended Data Figure 7 (Figure S7).**
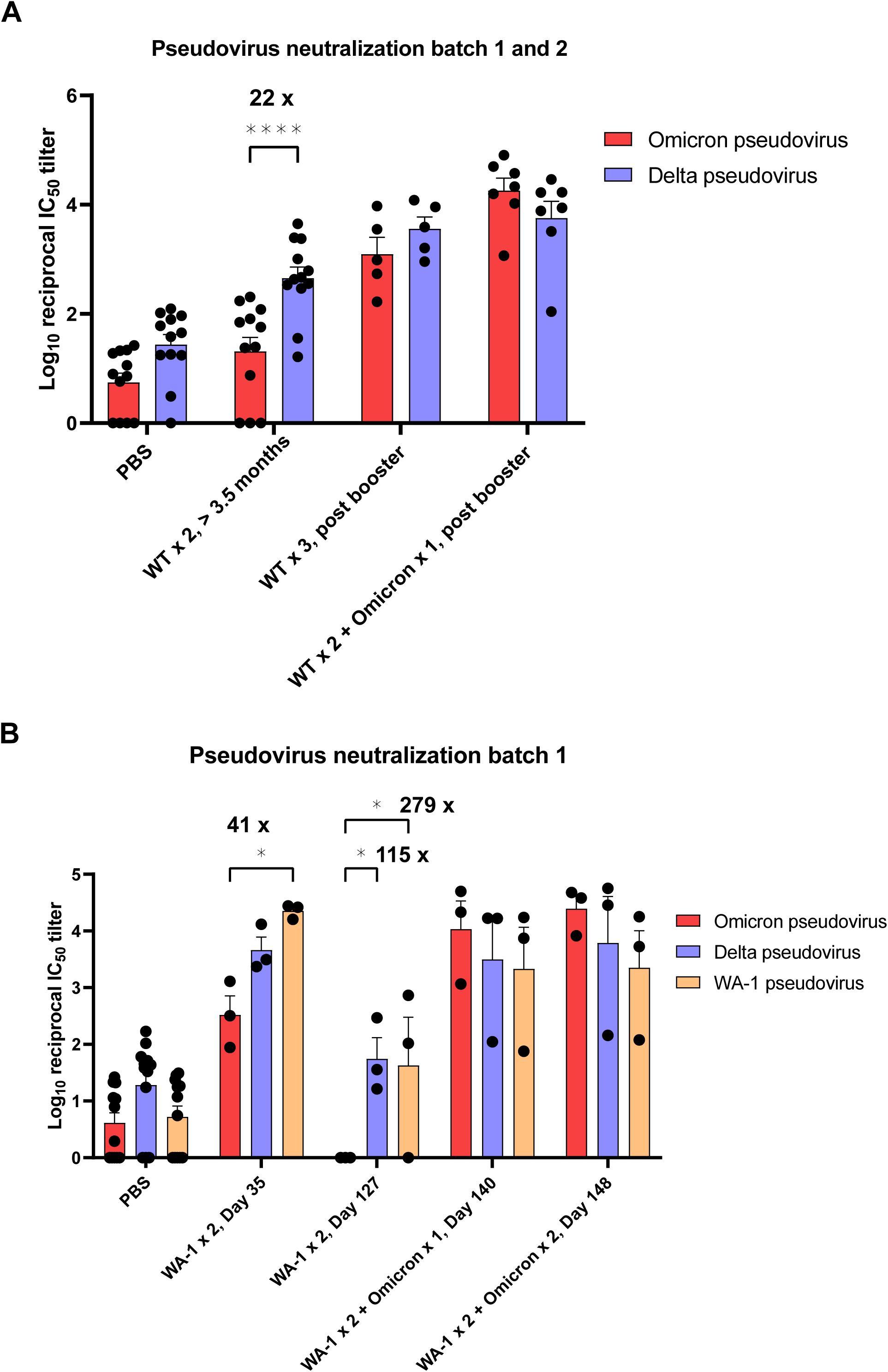
Neutralizing antibody titers in the form of log_10_-transformed reciprocal IC50 were grouped by vaccination timepoints to compare titers against different pseudoviruses. The neutralization titers from combined datasets (A) or batch 1 (B) were quantified as log_10_-transformed reciprocal IC50 values (Log_10_ reciprocal IC50, or Log_10_ IC50) based on titration curves in Extended Data Figure 6. Titer ratios were indicated in each graph and fold change is calculated from (ratio - 1). Data on dot-bar plots are shown as mean ± s.e.m. with individual data points in plots. Two-way ANOVA with Tukey’s multiple comparisons test was used to assess statistical significance. Statistical significance labels: * p < 0.05; ** p < 0.01; *** p < 0.001; **** p < 0.0001. Non-significant comparisons are not shown, unless otherwise noted as n.s., not significant.

**Extended Data Figure 8 (Figure S8).**
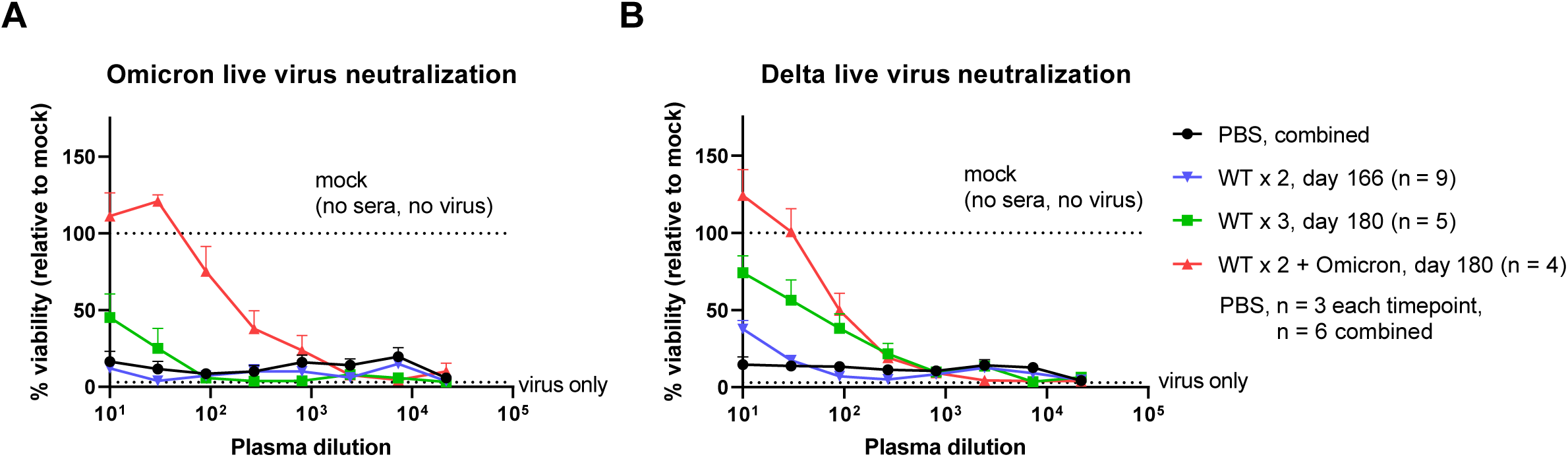
Live virus neutralization titration curves over serial dilution of plasma collected at different timepoints from mice administered with PBS or WT and/or Omicron LNP-mRNA. **A,** Omicron live virus titration curves (n = 9 before booster, n=5 in WT x 3, n = 4 in WT x 2 + Omicron) **B**. Delta live virus titration curves (n = 9 before booster, n=5 in WT x 3, n = 4 in WT x 2 + Omicron) Titration curves were plotted over serial dilution points of plasma collected from mice before and after WT or Omicron LNP-mRNA boosters at defined time points. Data of each sample were collected from two replicates.

**Extended Data Figure 9 (Figure S9).**
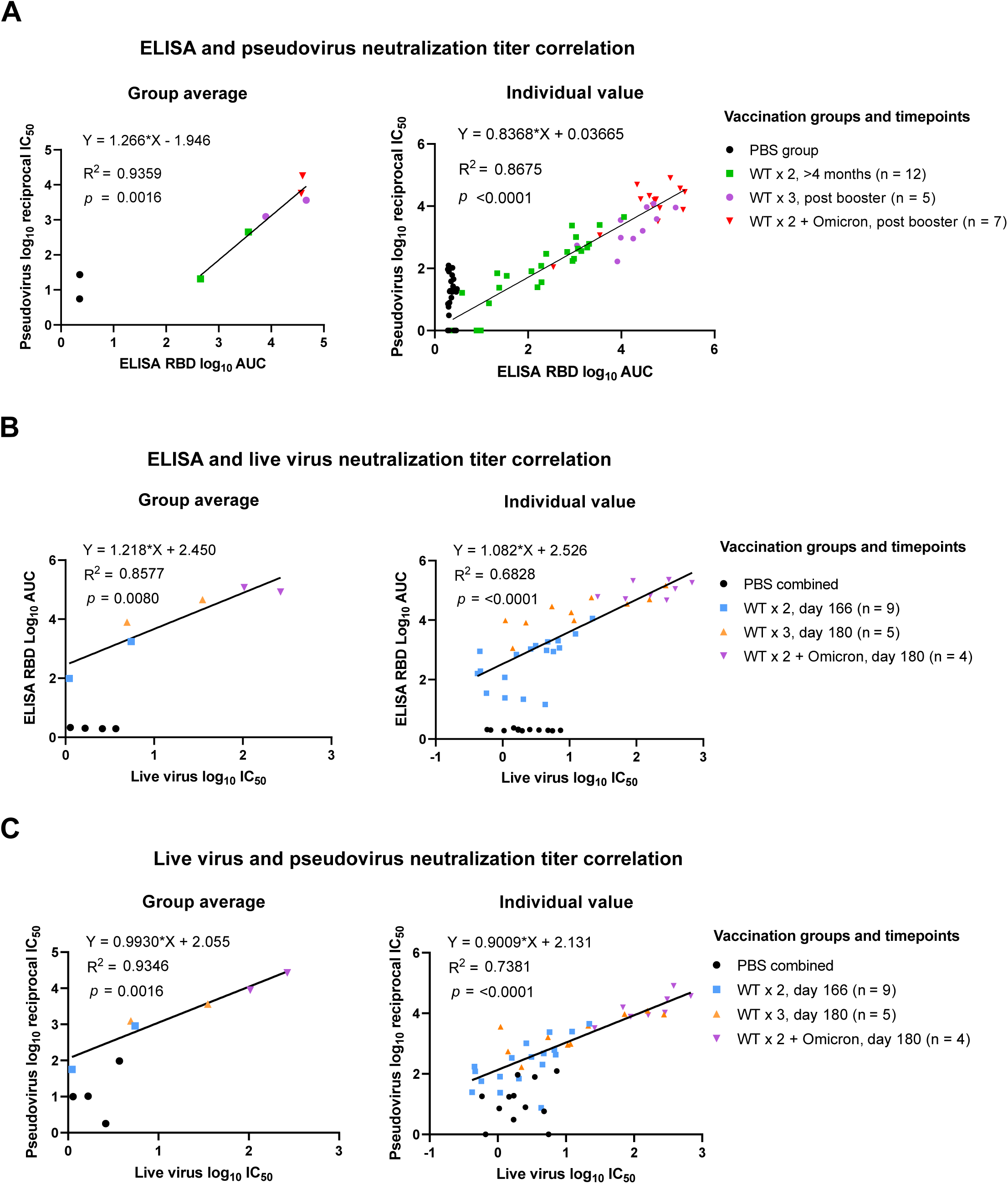
Correlation analysis of antibody titers determined by ELISA, pseudovirus neutralization and live virus neutralization assays. **A**, Correlation between pseudovirus neutralization titers (log_10_ reciprocal IC50, y axis) and ELISA titers (log_10_ AUC, x axis) from matched vaccination group (left panel) or individual mouse (right panel). **B**. Correlation between live virus neutralization titers (log_10_ IC50, x axis) and ELISA titers (log_10_ AUC, y axis) from matched vaccination group (left panel) or individual mouse (right panel). **C**. Correlation between live virus neutralization titers (log_10_ IC50, x axis) and pseudovirus neutralization titers (log_10_ AUC, y axis) from matched vaccination group (left panel) or individual mouse (right panel). PBS samples from different timepoints were shown as one group in correlation map and were not included in linear regression model. Each dot in bar graphs represents value from one group average (left panel), or one individual mouse (right panel).

**Extended Data Figure 10 (Figure S10).**
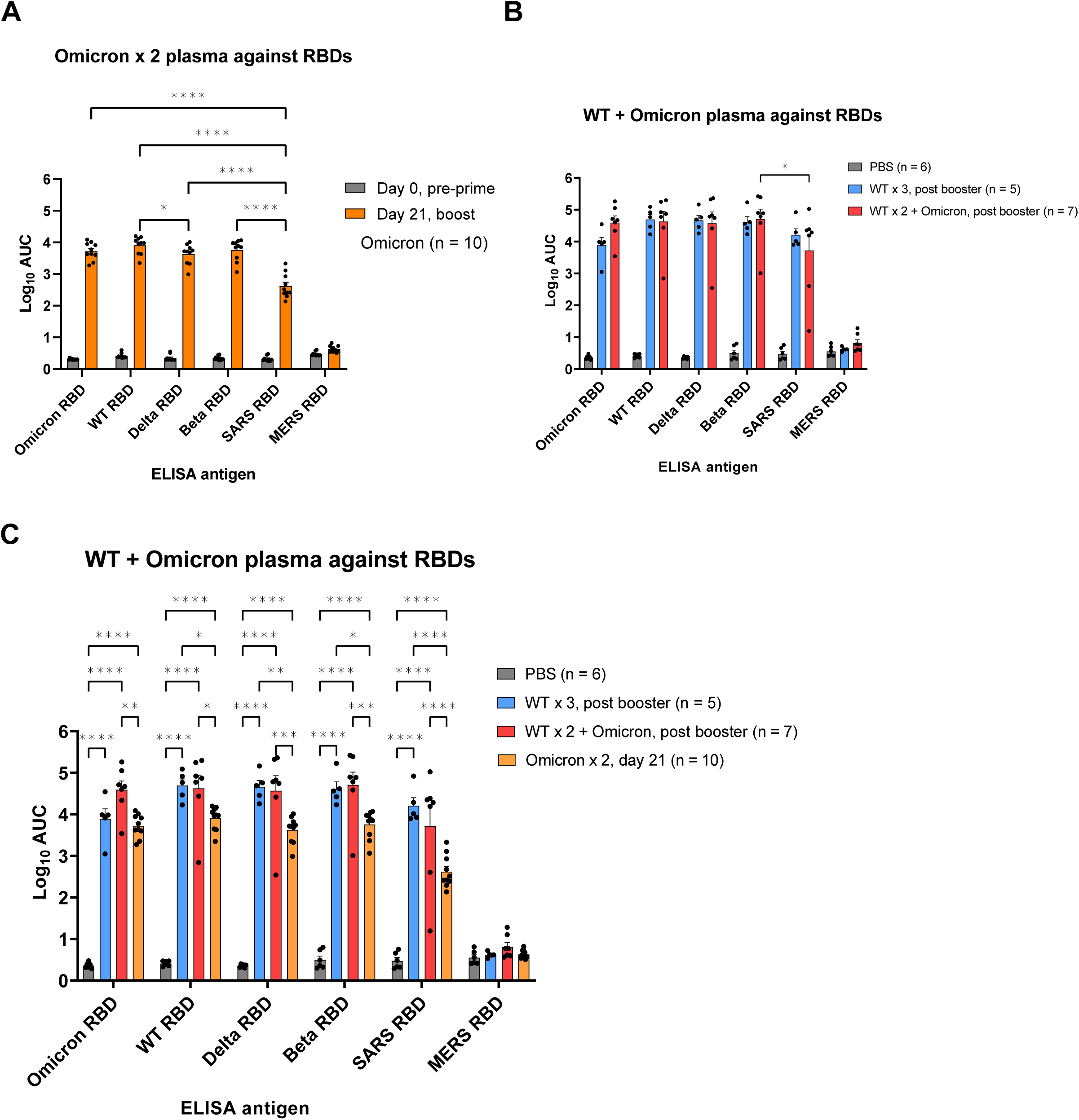
Assessment of WT or Omicron LNP-mRNA mediated cross reactivity against a panel of SARS-CoV-2 variants and pathogenic coronavirus species in ELISA. A, binding antibody titers (Log_10_ AUC) of plasma from mice that received Omicron LNP-mRNA prime and boost (Omicron x 2, n = 10). B, binding antibody titers of plasma from mice that received WT (WT x 3, n = 5) or Omicron (WT x 2 + Omicron, n = 7) LNP-mRNA boosters. This supplemental figure is a combination of data from the experiment shown in Figures 1 and 3 for comparison clarity. Data on dot-bar plots are shown as mean ± s.e.m. with individual data points in plots. Two-way ANOVA with Tukey’s multiple comparisons test was used to assess statistical significance. Multiple comparisons between titers against different ELISA antigens were made within same vaccination group. All comparisons with MERS RBD were significant and not shown in graph to simplify comparisons. Statistical significance labels: * p < 0.05; ** p < 0.01; *** p < 0.001; **** p < 0.0001. Non-significant comparisons are not shown.

**Extended Data Figure 11 (Figure S11).**
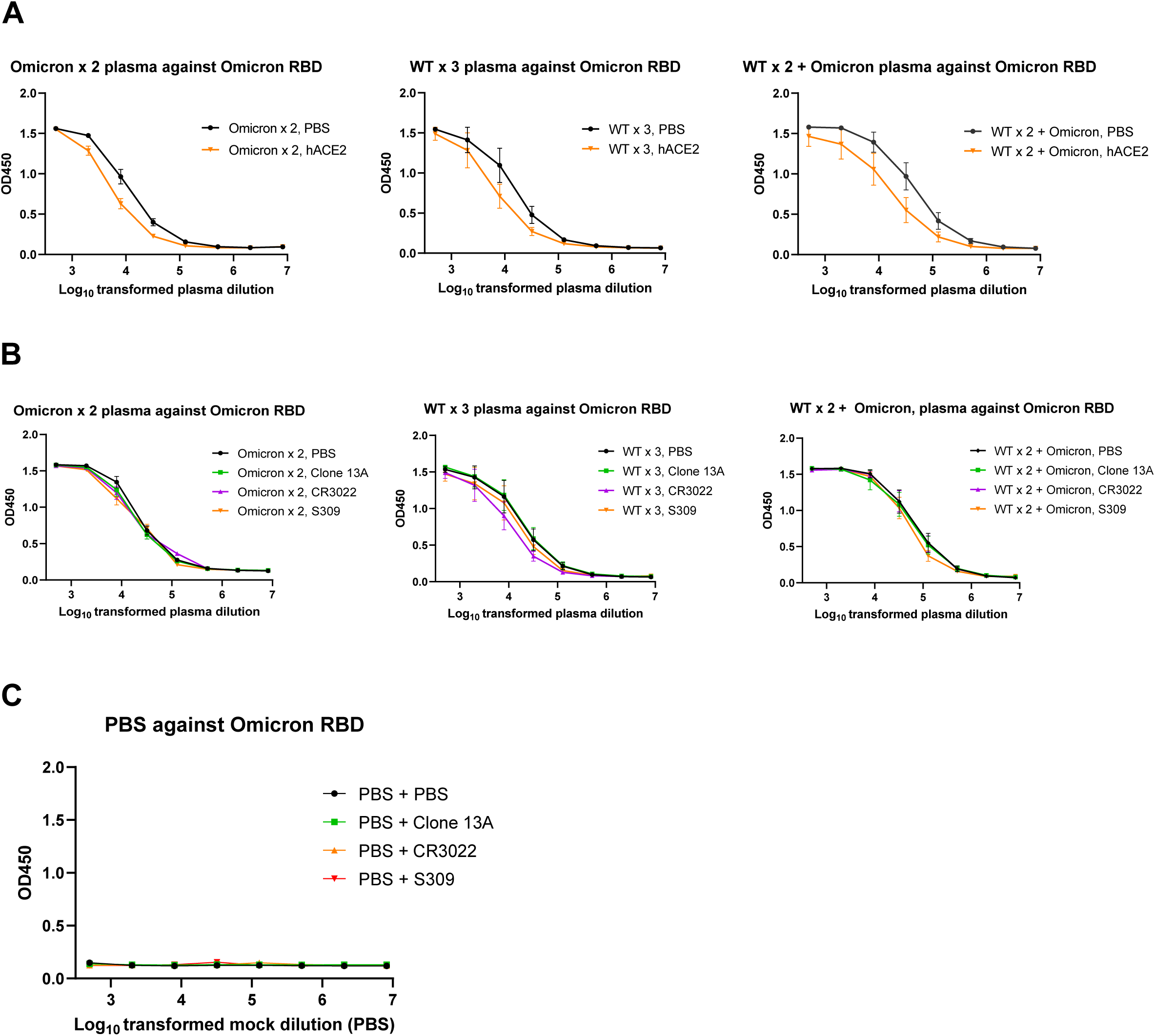
Competition ELISA titration curves and binding antibody titers against low-density Omicron RBD from mice vaccinated with WT and/or Omicron LNP-mRNA. **A.** hACE competition ELISA titration curves over a series of log_10_-transformed dilution points of plasma from mice vaccinated with Omicron LNP-mRNA (Omicron x 2 plasma, left, n = 10) or WT/Omicron LNP-mRNA (WT x 3, middle, n = 5 and WT x 2 + Omicron plasma, right, n = 7). **B**. antibody competition ELISA titration curves over a series of log_10_-transformed dilution points of plasma from mice vaccinated with Omicron LNP-mRNA (Omicron x 2 plasma, left, n = 10) or WT/Omicron LNP-mRNA (WT x 3 plasma, middle, n = 5 and WT x 2 + Omicron plasma, right, n = 7). **C**. PBS buffer as negative control to show minimal cross reactivity of anti-mouse secondary antibody with human IgG blocking antibodies, including Clone 13A, CR3022 and S309. n = 2 and each contains 8 mock dilution points.

**Extended Data Figure 12 (Figure S12).**
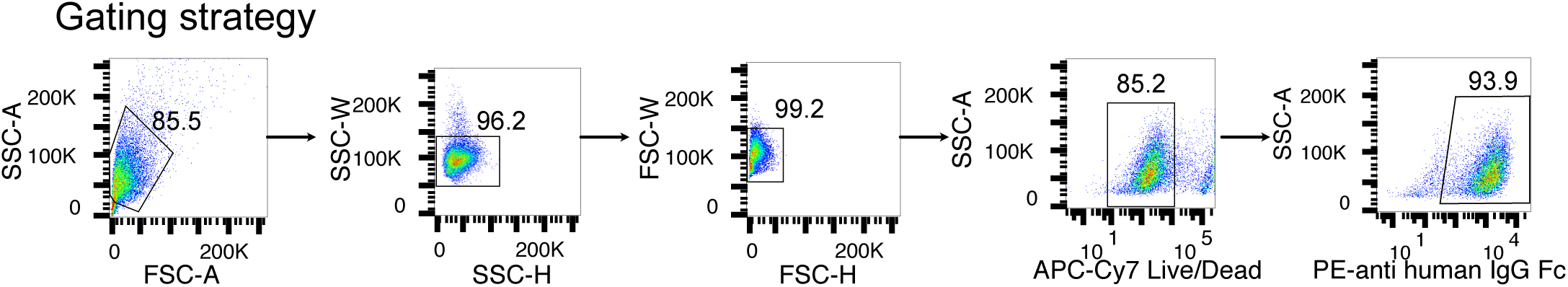
Representative flow cytometry gating strategy for detecting Omicron spike binding to human ACE2 receptor.

## Notes

### Competing Interest Statement

The authors have declared no competing interest.

### Summary of Updates

-(1) Performed parallel WT- and Omicron-specific vaccine booster comparison, to test the effect of homologous booster (WT x2+1) and heterologous booster (WT x2 + Omicron x1). Results showed that both WT- and Omicron-specific vaccine boosters enhanced the waning immunity, however, the effect of homologous Omicron-specific vaccine booster is stronger against the Omicron variant. -(2) Omicron vaccine vs Delta: Omicron-specific vaccine booster has comparable antibody response against Delta variant, i.e. it does not lose potency against Delta as compared to WT vaccine. -(3) Performed BL3 live virus (authentic virus) neutralization experiment, further validating the findings above. -(4) Performed broad antibody activity testing against VoCs (WT, Beta, Delta) and different Betacoronavirus species (SARS2, SARS, MERS), revealing that both Omicron alone and WT + Omicron vaccination showed broadly reactive antibody responses. -(5) Performed antibody characterization experiment in the context of other known clinical mAbs, showing that Omicron-mRNA induced antibody pool covering portions of antibodies contains Class I-III (overlapped with hACE2 epitopes), IV antibodies. -(6) Performed repeat experiments that increased sample sizes, e.g. numbers of animals used, experimental replicates, and improved statistics.

